# Inhibition of Hippo signalling disrupts germinative cell-cycle progression and larval development in *Echinococcus multilocularis*

**DOI:** 10.64898/2026.10.29.762824

**Authors:** Luisa Schiegl, Monika Bergmann, Klaus Brehm

## Abstract

The metacestode larval stage of *Echinococcus multilocularis* exhibits a remarkable capacity for sustained, infiltrative growth, driven by germinative cells, the parasite’s only mitotically active cell population. Yet, despite this seemingly unlimited growth potential, cell proliferation and differentiation must remain tightly coordinated to maintain tissue organization and parasite development. Here, we identify and functionally investigate components of the Hippo signaling pathway, a conserved regulator of cellular growth and tissue homeostasis, in *E. multilocularis*. We identified the principal components of the canonical pathway, including a highly divergent Yorkie ortholog, and demonstrated an interaction between EmYorkie and the TEAD-family transcription factor EmScalloped. Both *emhippo* and *emyorkie* are expressed in proliferating germinative cells and in association with differentiated neural and muscle cells, suggesting functions beyond stem cell regulation. RNA interference targeting *emyorkie* enhanced metacestode vesicle formation without a detectable increase in EdU incorporation at the time point examined. Treatment with XMU-MP-1, an inhibitor originally developed against mammalian MST1/2 kinases, induced pronounced enlargement and multinucleation of germinative cells, some of which retained the capacity for DNA synthesis despite the absence of detectable phospho-histone H3 staining. Comparison with the Aurora kinase inhibitor MLN8237 revealed differences in the magnitude and concentration dependence of giant-cell formation and phosphor-histone H3 staining. Together, our findings provide evidence for a role of EmYorkie in early larval development and reveal pronounced effects of XMU-MP-1 on germinative-cell morphology and cell-cycle-associated processes, highlighting the potential importance of Hippo-associated signaling in this parasitic flatworm.

## Introduction

*Echinococcus multilocularis*, the causative agent of alveolar echinococcosis (AE), is a medically important cestode parasite whose metacestode larval stage is characterized by extensive and potentially indefinite growth within the tissues of its intermediate hosts (Brunetti et al., 2010; Brehm and Koziol, 2017; Wen et al., 2019). In humans, the metacestode develops predominantly in the liver, where it proliferates by infiltrative growth and progressively affects surrounding tissues. AE is a severe and potentially fatal disease that is difficult to treat, particularly when complete surgical removal of the parasite is not possible (Brunetti et al., 2010). Chemotherapy relies mainly on benzimidazoles, particularly albendazole, which generally suppress parasite growth rather than reliably eliminating the infection. Consequently, patients with inoperable disease frequently require prolonged or lifelong treatment (Brunetti et al., 2010; Wen et al., 2019). The remarkable proliferative capacity of the metacestode, together with its resistance to curative chemotherapy, makes understanding the molecular regulation of parasite growth and development particularly important.

The continuous growth of the metacestode is sustained by a population of proliferating germinative cells (GC), which represent the only mitotically active cells of the parasite and produce all differentiated cells (Koziol et al., 2014). The GC support tissue renewal, vesicle expansion, and the formation of new parasite structures, including brood capsules and protoscoleces. The ability of GC to sustain long-term parasite growth while continuously generating differentiated progeny implies the existence of regulatory mechanisms that balance stem cell proliferation, differentiation, and tissue homeostasis. Their developmental potential is regulated by conserved signaling systems that also control stem cell behavior and tissue organization in other metazoans (Brehm and Koziol, 2017). Among these, WNT signaling has emerged as a major regulator of developmental patterning in *E. multilocularis*. WNT ligands expressed by muscle cells provide positional information, and interference with canonical WNT/β-catenin signaling profoundly alters GC behavior and metacestode development (Koziol et al., 2016; Herz et al., 2024; Herrmann et al., 2026). Thus, despite its unusual growth characteristics, the metacestode retains signaling mechanisms involved in coordinating cell proliferation, differentiation, and developmental organization.

An intriguing question is how the potentially unlimited growth of the metacestode relates to mechanisms that normally restrict tissue and organ size in other animals. Whereas the oncosphere, protoscolex, and adult worm develop characteristic body dimensions and defined anatomical structures, the metacestode lacks an obvious predetermined final size and can continue expanding throughout the life of its host. This raises the possibility that conserved growth-control mechanisms have acquired modified functions during metacestode development or that their regulatory activity differs between parasite life-cycle stages. One of the most prominent signaling systems involved in the regulation of tissue growth and organ size in metazoans is the Hippo pathway.

Originally characterized in *Drosophila melanogaster*, the Hippo pathway regulates cell proliferation, apoptosis, stem cell activity, and tissue homeostasis through a conserved protein kinase cascade (Harvey and Tapon, 2007; Pan, 2010; Yu et al., 2015). In its canonical configuration, the kinase Hippo (MST1/2 in mammals), together with its regulatory partner Salvador (SAV1), activates the downstream kinase Warts (LATS1/2) and its cofactor Mats (MOB1). Warts/LATS-mediated phosphorylation promotes the cytoplasmic retention and inactivation of the transcriptional coactivator Yorkie (YAP/TAZ in mammals), thereby restricting its nuclear activity in association with the transcription factor Scalloped (TEAD in mammals). Beyond its classical role in organ-size control, Hippo signaling also influences cell fate decisions, cytoskeletal organization, and interactions with other developmental pathways, including WNT/β-catenin signaling (Piccolo et al., 2014; Misra and Irvine, 2018). Studies in planarians have demonstrated that Yorkie/Hippo signalling regulates stem cell proliferation, cell-cycle progression, tissue homeostasis, and regenerative patterning (Lin and Pearson, 2014, 2017; de Sousa et al., 2018). These findings suggest that Hippo signalling has important functions in coordinating stem cell activity and developmental organization in flatworms. However, the organization and biological functions of this pathway remain largely unexplored in parasitic flatworms.

In the present study, we investigated the Hippo signaling pathway in *E. multilocularis*, focusing on its potential involvement in metacestode development and GC regulation. We identified the principal components of the conserved Hippo signaling machinery, characterized a divergent Yorkie ortholog and its interaction with the TEAD-family transcription factor Scalloped, and examined the expression of Hippo pathway components across larval stages and cellular populations. We further investigated the effects of RNAi-mediated interference with *emhippo* and *emyorkie* on metacestode vesicle formation and analyzed the consequences of treatment with the MST1/2 inhibitor XMU-MP-1 for GC morphology, cell-cycle progression, and parasite tissue integrity. Together, these approaches allowed us to investigate the contribution of Hippo-associated signalling to GC regulation, cell-cycle progression, and developmental organization in a parasitic flatworm characterized by sustained and seemingly unrestricted larval growth.

## Materials and Methods

### Parasite material and *in vitro* cultivation

The *E. multilocularis* isolates H95, GH09, and KB19 were used in this study. Isolate H95 originated from a naturally infected red fox (Vulpes vulpes) in Germany (Jura et al., 1996), whereas GH09 and KB19 were derived from naturally infected Old World monkeys (*Macaca spp.*) housed in a breeding facility in Göttingen, Germany (Tappe et al., 2007). Metacestode material was maintained by serial intraperitoneal passage in Mongolian jirds (*Meriones unguiculatus*) and cultivated in vitro according to established procedures (Spiliotis et al., 2004; Spiliotis and Brehm, 2009).

For in vitro cultivation, parasite material was co-cultured with rat Reuber hepatoma (RH−) cells (ATCC CRL-1600) in Dulbecco’s modified Eagle’s medium (DMEM) supplemented with 10% fetal bovine serum and 1% penicillin/streptomycin. Cultures were maintained at 37°C in a humidified atmosphere containing 5% CO₂. Metacestode vesicles were subsequently transferred to axenic conditions by repeated washing with phosphate-buffered saline (PBS) and cultivation in RH− cell-conditioned medium supplemented with reducing agents under a nitrogen atmosphere, as previously described (Spiliotis et al., 2004; Spiliotis and Brehm, 2009). Vesicles were maintained under axenic conditions for at least three days before experimental manipulation. Primary cells were isolated from axenically cultivated metacestode vesicles of isolates H95 and GH09 by mechanical disruption and trypsin-mediated dissociation, followed by differential centrifugation and filtration as described by Spiliotis and Brehm (2009) and Spiliotis et al. (2010). Cell suspensions were cultured in a 1:1 mixture of A6 and B4 RH− cell-conditioned media under oxygen-restricted conditions at 37°C. Culture medium was renewed every two to three days. Protoscoleces were isolated from infected jird tissue by mechanical dissociation, sieving, and repeated washing with PBS. Activation was performed by incubation in DMEM containing 0.05% pepsin at pH 2.0 for 20 min at 37°C, followed by washing and incubation in DMEM containing 0.2% sodium taurocholate for 3 h at 37°C. Activated protoscoleces were maintained in conditioned medium under a nitrogen atmosphere.

### Identification and sequence analysis of Hippo pathway components

Putative components of the *E. multilocularis* Hippo signaling pathway were identified by homology-based searches using protein sequences from *Drosophila melanogaster*, *Schmidtea mediterranea*, and *Homo sapiens*. Reference sequences were retrieved from UniProt and used as queries in BLASTP searches against the *E. multilocularis* genome and predicted proteome available through WormBase ParaSite. Candidate sequences were subjected to reciprocal similarity searches against annotated protein databases to assess their evolutionary relationships. Predicted protein sequences were examined for conserved domains and characteristic sequence motifs using SMART (https://smart.embl.de/) and comparative sequence alignments. Because initial searches using *Drosophila* Yorkie did not identify an unequivocal ortholog, additional searches were performed using the more closely related Yorkie sequence from *S. mediterranea*. The predicted *emyorkie* transcript model (https://parasite.wormbase.org) was examined against available sequencing evidence and corrected where necessary.

### Yeast two-hybrid analysis

Interactions between EmYorkie and EmScalloped were investigated using the Matchmaker GAL4 yeast two-hybrid system (Clontech). Coding sequences were cloned into the GAL4 activation-domain vector pGADT7 and DNA-binding-domain vector pGBKT7. Both bait–prey orientations were examined. Primer sequences for plasmid construction are listed in Supplementary Table 1. The resulting constructs were co-transformed into *Saccharomyces cerevisiae* strain AH109. Transformants were initially selected on synthetic dropout medium lacking leucine and tryptophan (SD/−Leu/−Trp). Protein interactions were subsequently assessed by growth on SD/−Leu/−Trp/−His and SD/−Leu/−Trp/−His/−Ade media, representing intermediate and high selection stringency, respectively. The interaction between SV40 large T-antigen and p53 served as a positive control, while lamin C/T-antigen combinations and constructs paired with the corresponding empty vectors served as negative controls. Interactions were examined in three independent co-transformation experiments, using individual colonies and pooled transformants for growth assays.

### RNA interference and quantitative RT-PCR

RNA interference was performed in *E. multilocularis* primary cell cultures using synthetic small interfering RNAs (siRNAs) targeting *emhippo* or *emyorkie*, individually or in combination. An siRNA targeting *gfp*, which has no corresponding endogenous sequence in *E. multilocularis*, served as a negative control (Herrmann et al., 2026). Candidate siRNA sequences were selected using the Thermo Fisher RNAi Designer and screened against the predicted parasite transcriptome to minimize potential off-target effects. siRNA sequences used are listed in Supplementary Table 1. Primary cell aggregates were subjected to electroporation following the approach established by Spiliotis et al. (2010). For each electroporation, 180 primary cell units were resuspended in 90 µl electroporation buffer containing 2 µl of a 100 µM siRNA solution. Electroporation was performed in 1-mm cuvettes at 200 V for 0.6 ms. Following a 12-min recovery period at 37°C, cells were transferred to conditioned medium and seeded into 96-well plates. Cultures were maintained for seven days, with medium replacement every two to three days. Vesicle formation was assessed microscopically. For proliferation analysis, primary cell aggregates were labeled with EdU on day 7. Additional samples were collected on days 2 and 7 for RNA isolation and analysis of knockdown efficiency.

Total RNA was isolated using the Direct-zol RNA Miniprep Kit (Zymo Research), including DNase treatment. Complementary DNA was synthesized using SuperScript IV reverse transcriptase (Thermo Fisher Scientific). Quantitative PCR was performed using SYBR Green chemistry with gene-specific primers. Amplification efficiencies were determined from standard dilution series, and specificity was assessed by melting-curve analysis. Transcript levels were normalized against the reference gene *emelp*, and relative expression was calculated using the comparative 2^−ΔΔCt method (Livak and Schmittgen, 2001).

### Analysis of transcriptomic datasets

Expression of *emhippo* and *emyorkie* was examined using previously generated transcriptomic datasets covering different developmental stages and experimental conditions of *E. multilocularis*. These included primary cell cultures, metacestode vesicles, activated and non-activated protoscoleces, hydroxyurea-treated metacestode tissue, and samples subjected to β-catenin knockdown (Tsai et al., 2013; Herz et al., 2024; Herrmann et al., 2026). Expression values were compared across the respective datasets to evaluate developmental regulation and potential enrichment in proliferating or differentiated cell populations.

### Whole-mount in situ hybridization

Whole-mount in situ hybridization (WISH) was performed on metacestode germinal-layer preparations and activated protoscoleces using digoxigenin-labeled RNA probes, following established procedures for *E. multilocularis* (Koziol et al., 2014; Koziol et al., 2016; Herrmann et al., 2026). Gene-specific fragments were amplified and cloned into plasmid vectors, and antisense RNA probes were synthesized by in vitro transcription using T7 or SP6 RNA polymerase. Fixed specimens were rehydrated, permeabilized with proteinase K, acetylated with acetic anhydride, and post-fixed in 4% paraformaldehyde. After prehybridization, specimens were incubated overnight with the respective RNA probes at 58°C. Following stringent post-hybridization washes, probes were detected using peroxidase-conjugated anti-digoxigenin antibodies and fluorescein-tyramide signal amplification. Hybridized specimens were subsequently processed for EdU detection, immunofluorescence, or nuclear staining, as required.

### EdU incorporation and immunofluorescence

DNA synthesis was assessed by incorporation of 5-ethynyl-2′-deoxyuridine (EdU) as previously described (Koziol et al., 2014; Herz et al., 2024). Unless otherwise indicated, metacestode vesicles and primary cell cultures were exposed to 50 µM EdU for 5 h at 37°C before fixation. Protoscoleces were labeled under the conditions specified for the respective experiments. Following fixation in 4% paraformaldehyde, EdU was detected using the Click-iT EdU Alexa Fluor 555 Imaging Kit (Thermo Fisher Scientific) according to the manufacturer’s instructions. Cell nuclei were counterstained with DAPI. For immunofluorescence, fixed samples were permeabilized, blocked with bovine serum albumin and sheep serum, and incubated with primary antibodies overnight at 4°C. After extensive washing, specimens were incubated with fluorophore-conjugated secondary antibodies. Antibodies against acetylated α-tubulin (Koziol et al., 2013) and high-molecular-weight tropomyosin (Koziol et al., 2016) were used to visualize neuronal and muscle-associated structures, respectively. The germinative-cell-associated β-tubulin isoform Tub-2 was detected using a specific antibody (Bergmann et al., 2026), while phosphorylated histone H3 (pH3) was used as a marker of mitotic cells (Koziol et al., 2014).

### Pharmacological treatments

The effects of XMU-MP-1, an inhibitor originally developed against mammalian MST1/2 kinases, were examined in axenically cultivated metacestode vesicles, primary cell cultures, and activated protoscoleces. XMU-MP-1 (Selleckchem) was dissolved in dimethyl sulfoxide (DMSO) to prepare a 10 mM stock solution and applied at final concentrations of 1 or 10 µM. Corresponding DMSO-treated cultures served as vehicle controls. For metacestode experiments, intact vesicles approximately 3–5 mm in diameter were distributed into 12-well plates, generally with 10–11 vesicles per well. After an adaptation period, cultures were exposed to inhibitor-containing conditioned medium under a nitrogen atmosphere at 37°C. Medium containing freshly diluted inhibitor was replaced every two to three days. Vesicle integrity and morphological alterations were monitored microscopically throughout treatment. Primary cell cultures were treated with 1 or 10 µM XMU-MP-1 for up to 14 days. Activated protoscoleces were exposed to the same concentrations, with medium changes every two to three days. At the indicated time points, specimens were processed for microscopy, EdU incorporation, WISH, or immunofluorescence. To compare the effects of XMU-MP-1 with those of Aurora kinase inhibition, metacestode vesicles were treated with XMU-MP-1 or MLN8237 (Alisertib; MedChem) at concentrations of 1 and 5 µM. The number of pH3-positive cells was determined after 24 h and seven days of treatment. Giant-cell formation and EdU incorporation were analyzed in parallel experiments.

### Microscopy, image analysis, and quantification

Fluorescence microscopy was used to assess gene expression, EdU incorporation, immunostaining, and cellular morphology. Images were analyzed using ImageJ. For WISH/EdU experiments, the proportions of transcript-positive and EdU-positive cells were determined from defined germinal-layer regions. Giant cells were identified according to their enlarged cellular morphology and quantified as the number of cells per unit area of germinal-layer tissue. Mitotic activity was assessed by counting pH3-positive cells and expressing the results as cells per mm². Where applicable, EdU incorporation was quantified separately in giant cells and cells of regular morphology. Metacestode vesicle integrity was assessed by microscopic inspection, and vesicles exhibiting structural collapse were recorded at the indicated time points.

### Statistical analysis

Statistical analyses were performed using GraphPad Prism. Data are presented as mean ± standard deviation unless otherwise indicated. Experiments were performed with independent biological replicates, and the number of replicates is specified in the corresponding figure legends. Comparisons involving multiple experimental groups were analyzed using one-way or two-way analysis of variance (ANOVA), as appropriate, followed by Dunnett’s multiple-comparison test against the corresponding control. Differences were considered statistically significant at p < 0.05.

## Results

### Identification of conserved Hippo pathway components in *E. multilocularis*

To investigate the presence of a Hippo signaling pathway in *E. multilocularis*, we performed homology-based searches using known pathway components from *D. melanogaster* and humans as queries (Ma et al., 2019). These analyses identified putative orthologs of the central components of the canonical Hippo signaling cascade, including the serine/threonine kinases Hippo (EmuJ_000067500) and Warts (EmuJ_000700100), their regulatory cofactors Salvador (EmuJ_000331400) and Mats (EmuJ_000189500), as well as the transcriptional coactivator Yorkie (EmuJ_000808000) and the TEAD-family transcription factor Scalloped (EmuJ_000606300). For each of these components, only a single candidate gene was identified in the *E. multilocularis* genome, with no evidence for additional paralogs. The proposed organization of the pathway in *E. multilocularis* is illustrated in Fig. 1.

**Figure 1.**
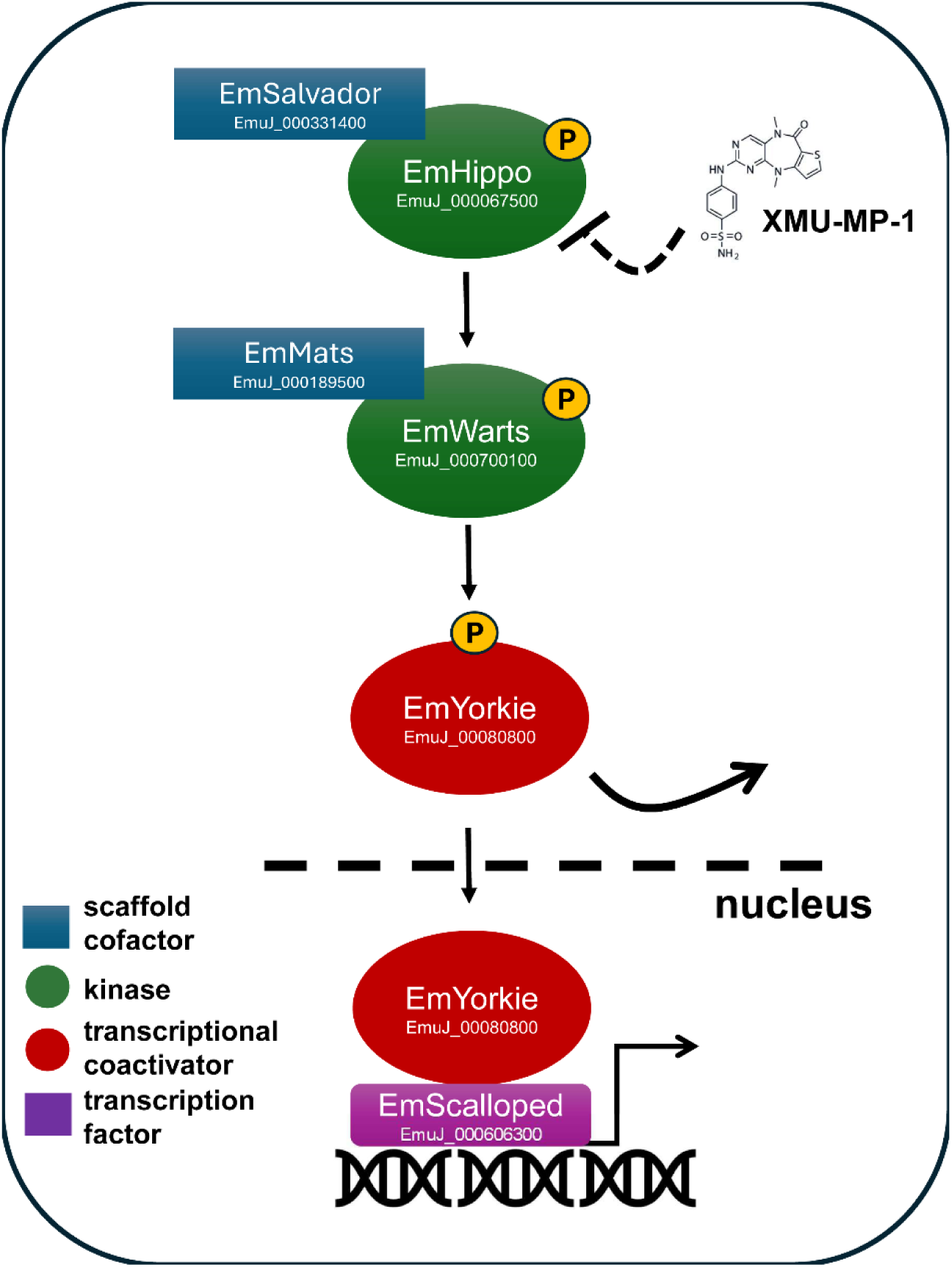
Schematic representation of the canonical Hippo signaling pathway in *E. multilocularis*. The proposed Hippo signaling cascade comprises the conserved serine/threonine kinases EmHippo and EmWarts, together with their respective regulatory cofactors EmSalvador and EmMats. In the canonical pathway, Hippo activates Warts through phosphorylation, which in turn phosphorylates the transcriptional coactivator EmYorkie, thereby preventing its nuclear activity. In the absence of inhibitory phosphorylation, EmYorkie can translocate into the nucleus, where it interacts with the TEAD-family transcription factor EmScalloped to regulate target gene expression. The small-molecule inhibitor XMU-MP-1, an established inhibitor of mammalian MST1/2 kinases, is proposed to interfere with Hippo kinase activity in *E. multilocularis*. The corresponding *E. multilocularis* gene identifiers (EmuJ) are indicated. The diagram represents a model based on the conserved canonical Hippo pathway; individual phosphorylation events and direct inhibition of EmHippo by XMU-MP-1 have not yet been experimentally verified in the parasite.

Comparative sequence analyses revealed conservation of characteristic functional domains among the identified proteins. EmHippo contains a conserved N-terminal serine/threonine kinase domain and a C-terminal SARAH domain, characteristic of Hippo/MST family kinases (Supplementary Fig. S1). Notably, several amino acid residues involved in the binding of XMU-MP-1 to mammalian MST2, as identified by crystallographic analysis (Fan et al., 2016), are conserved in EmHippo (Supplementary Fig. S1). This suggests that EmHippo represents a potential target of XMU-MP-1 in *E. multilocularis*. EmWarts contains a conserved serine/threonine kinase domain and a characteristic AGC kinase C-terminal region (Supplementary Fig. S3). The regulatory proteins EmMats and EmSalvador also exhibit conserved sequence features characteristic of their respective protein families (Supplementary Figs. S2 and S4). Similarly, EmScalloped contains a highly conserved TEA DNA-binding domain and a C-terminal region showing homology to other TEAD-family transcription factors (Supplementary Fig. S5).

Together, these findings indicate that *E. multilocularis* possesses the principal molecular components required for a canonical Hippo signaling cascade, suggesting that the basic architecture of this pathway has been conserved during flatworm evolution.

### Identification of a divergent Yorkie ortholog and its interaction with EmScalloped

While the core kinase components of the Hippo signaling pathway could be readily identified in the *E. multilocularis* genome, identification of the downstream transcriptional coactivator Yorkie proved more challenging. Initial homology searches using the *D. melanogaster* Yorkie sequence failed to identify an unambiguous ortholog, yielding predominantly candidates with similarities to ubiquitin ligases. We therefore extended our searches to Yorkie proteins from evolutionarily more closely related flatworms. Using the Yorkie sequence from the planarian *Schmidtea mediterranea* (Lin and Pearson, 2014), we identified EmuJ_000808000 as a potential Yorkie ortholog. Following correction of the predicted transcript model, the deduced protein contained a single WW domain and showed limited but recognizable sequence similarity to vertebrate YAP proteins. Nevertheless, the relatively low overall sequence conservation did not allow an unequivocal assignment based on sequence comparisons alone.

To obtain additional evidence for the identity of this candidate, we investigated its interaction with EmScalloped, the *E. multilocularis* ortholog of the TEAD-family transcription factors. In contrast to Yorkie, EmScalloped could be readily identified by sequence homology, including the presence of a highly conserved TEA DNA-binding domain (Supplementary Fig. S5). Since the interaction between Yorkie/YAP and Scalloped/TEAD represents a conserved feature of canonical Hippo signaling (Ma et al., 2019), we examined whether the candidate EmYorkie protein interacted with EmScalloped in yeast two-hybrid assays. A strong interaction was detected in both bait–prey orientations, whereas the corresponding negative controls showed no detectable interaction (Fig. 2B). These findings provide experimental support for the assignment of EmuJ_000808000 as the *E. multilocularis* Yorkie ortholog.

**Figure 2.**
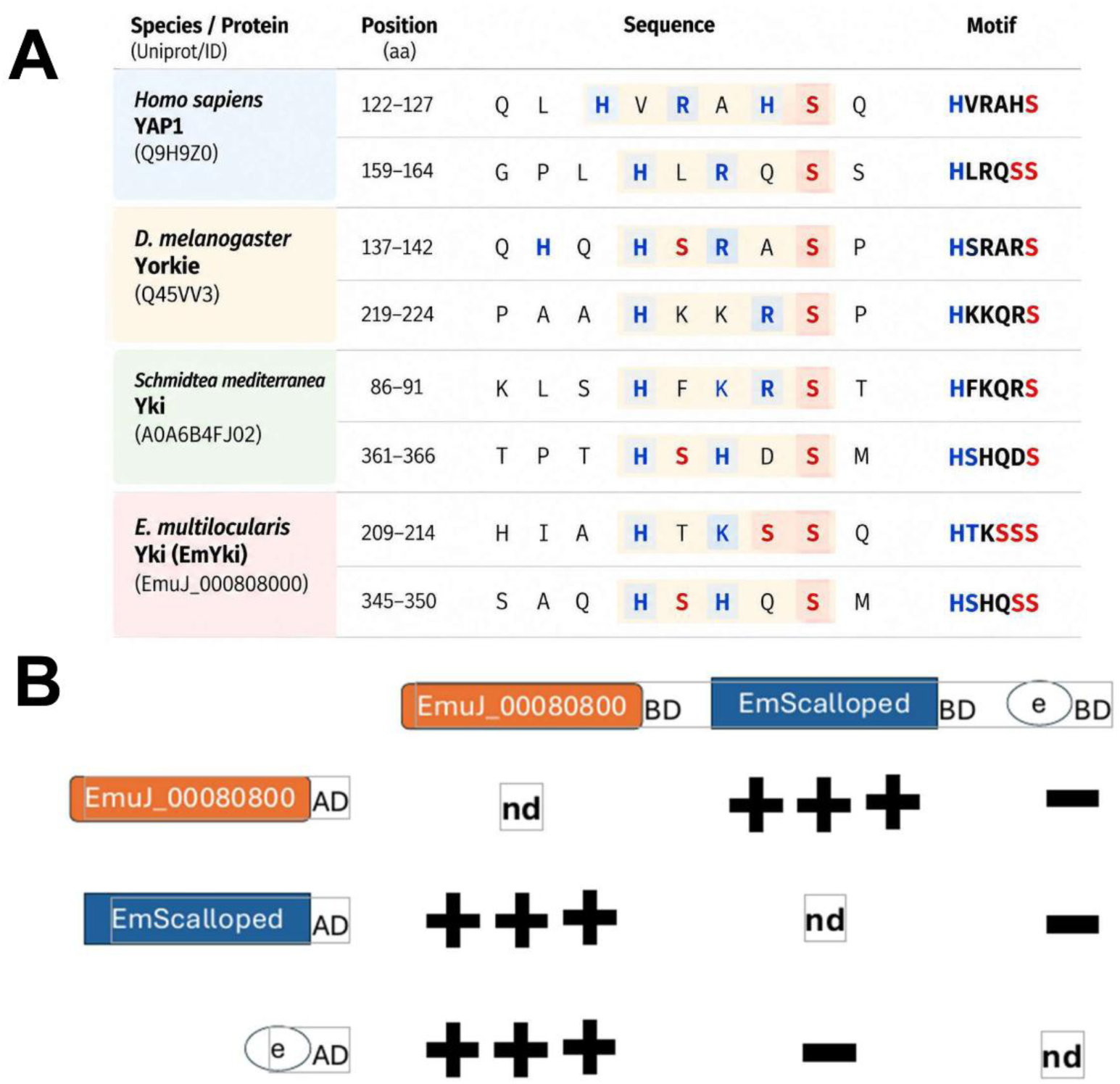
Putative Warts/LATS phosphorylation motifs in EmYorkie and interaction with EmScalloped. **(A)** Comparison of Warts/LATS phosphorylation motifs in human YAP1, *Drosophila melanogaster* Yorkie, *Schmidtea mediterranea* Yorkie, and *Echinococcus multilocularis* Yorkie (EmYorkie). Experimentally characterized phosphorylation sites in human YAP1 (S127 and S164; Zhao et al., 2007) and *D. melanogaster* Yorkie (S168 and S250; Oh and Irvine, 2009) are shown alongside corresponding candidate motifs in *S. mediterranea* Yorkie and EmYorkie. The six-amino-acid Warts/LATS recognition motif, Hx(R/K/H)xx(S/T), is highlighted, with three additional amino acids shown on either side. Histidine and basic residues are indicated in blue, whereas serine or threonine residues at the putative phosphorylation position are highlighted in red. Amino acid positions and protein accession numbers are indicated. Phosphorylation of the candidate sites in *S. mediterranea* and *E. multilocularis* has not been experimentally demonstrated. **(B)** Yeast two-hybrid analysis of the interaction between EmYorkie (EmuJ_000808000) and EmScalloped. Full-length coding sequences were expressed as fusion proteins with the GAL4 activation domain (AD) or DNA-binding domain (BD), as indicated. Empty vectors (e) served as negative controls. Interactions were assessed by yeast growth on selective media. Growth on SD/−Leu/−Trp/−His was considered indicative of a weak interaction, whereas growth on SD/−Leu/−Trp/−His/−Ade indicated a strong interaction (+++). No detectable interaction is indicated by (−), and nd denotes not determined. Three independent yeast two-hybrid experiments were performed, examining nine individual colonies and one pooled sample per interaction pair.

We further examined whether EmYorkie retained potential regulatory motifs characteristic of Yorkie/YAP proteins. Although the overall amino acid sequence conservation was low, sequence comparisons with human YAP1 and Yorkie proteins from *D. melanogaster* and *S. mediterranea* revealed two candidate Warts/LATS phosphorylation motifs in EmYorkie (Fig. 2A). These motifs resemble sequences associated with phosphorylation-dependent regulation of Yorkie/YAP in other organisms (Zhao et al., 2007; Oh and Irvine, 2009). Their presence provides additional support for the proposed identity of EmYorkie and suggests that the protein may be regulated by EmWarts.

### Expression of EmHippo during larval development

To investigate the potential involvement of Hippo signaling in larval development, we examined *emhippo* expression in publicly available transcriptomic datasets covering different developmental stages and experimental conditions of *E. multilocularis* (Tsai et al., 2013; Herz et al., 2024; Herrmann et al., 2026). *Emhippo* transcripts were detected across the investigated larval stages, including metacestodes and protoscoleces, as well as in primary cell cultures (Fig. 3B). We further examined datasets obtained from metacestodes subjected to hydroxyurea (HU) treatment, which depletes proliferating GC, and from metacestodes following β-catenin knockdown. Notably, *emhippo* transcript levels were not reduced following HU treatment and showed no significant changes upon β-catenin knockdown (Fig. 3B). These findings suggest that *emhippo* expression is not restricted to proliferating GC and that its overall transcript abundance is not significantly affected by β-catenin knockdown under the conditions examined.

**Figure 3.**
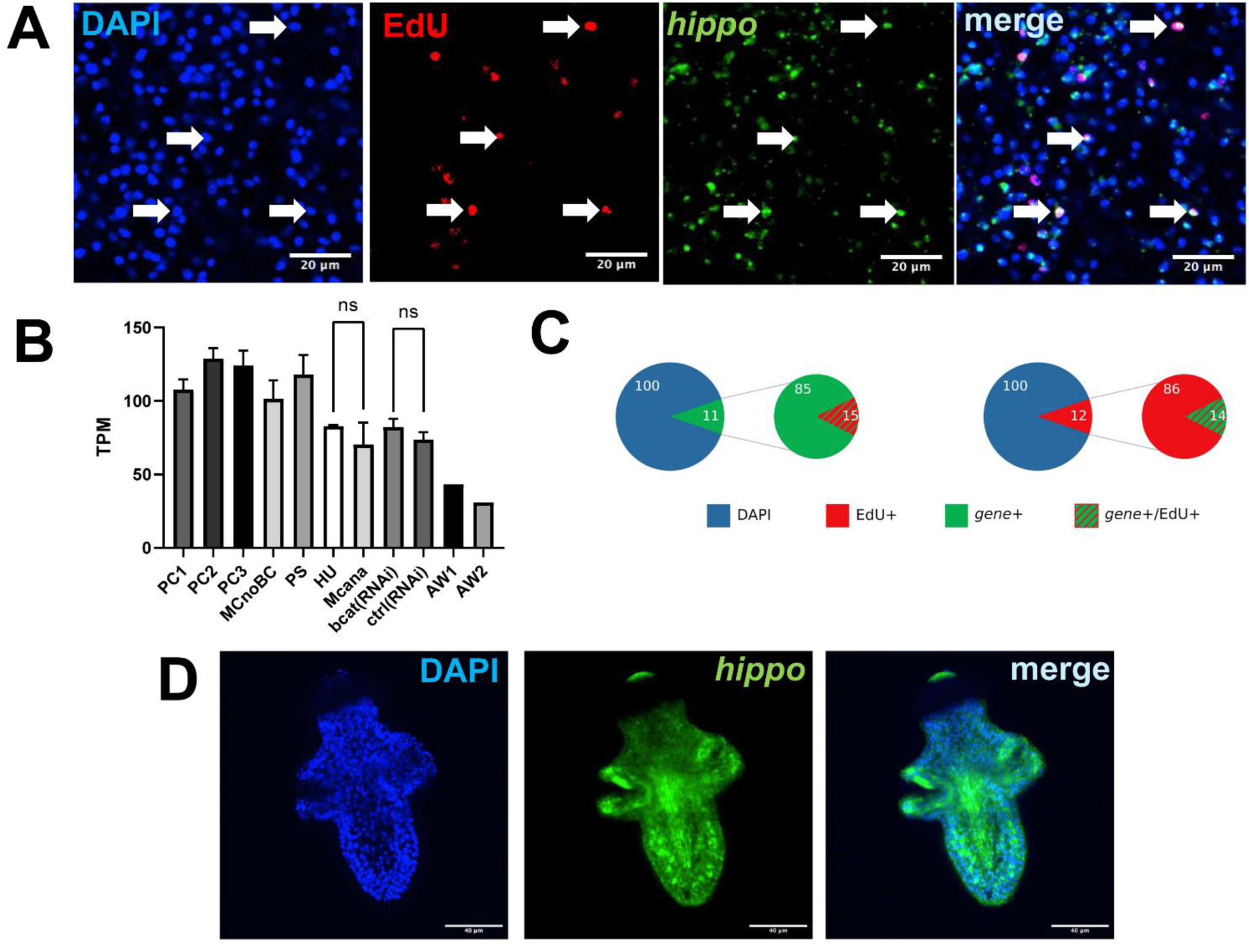
Expression of *emhippo* in *E. multilocularis* metacestode and protoscoleces. **(A)** WISH analysis of *emhippo* expression in the germinal layer of metacestode vesicles. *emhippo* transcripts are shown in green, EdU incorporation in red, and nuclei counterstained with DAPI in blue. Merged images illustrate the distribution of *emhippo*-expressing cells relative to proliferating cells. Arrows indicate representative EdU+ cells expressing *emhippo*. Scale bars: 20 µm. **(B)** Transcriptomic expression profile of *emhippo* across different developmental stages and experimental conditions, based on RNA-seq datasets analyzed as described by Tsai et al., 2013, Herz et al. 2024, and Herrmann et al., 2026. Expression levels are presented as transcripts per million (TPM). PC1–PC3, primary cell cultures; MCnoBC, metacestodes without brood capsules; PS, protoscoleces; HU, hydroxyurea-treated metacestodes; MCana, metacestodes maintained under anaerobic conditions; bcat(RNAi), β-catenin RNAi-treated samples; ctrl(RNAi), corresponding RNAi controls; AW1, pre-gravid adult worm; and AW2, gravid adult worm. Error bars indicate variation between samples. ns, not statistically significant. **(C)** Quantification of *emhippo* expression and EdU incorporation in the germinal layer of metacestode vesicles. The proportions of *emhippo*-positive and EdU-positive cells were determined relative to the total number of DAPI-stained nuclei (100%). The enlarged diagrams illustrate the fractions of marker-positive cells showing overlap between *emhippo* expression and EdU incorporation. Green, *emhippo*-positive cells; red, EdU-positive cells; hatched areas, double-positive cells. **(D)** WISH hybridization analysis of *emhippo* expression in activated protoscolex. *emhippo* transcripts are shown in green and DAPI-stained nuclei in blue. The merged image illustrates the broad distribution of *emhippo* expression throughout the protoscolex. Scale bars: 40 µm. All pictures represent single confocal slices.

We next investigated the spatial distribution of *emhippo* transcripts in the metacestode germinal layer using WISH, combined with EdU labeling to identify proliferating cells. *Emhippo*-expressing cells were distributed throughout the germinal layer without an obvious spatial clustering pattern (Fig. 3A). Quantitative analysis revealed that 12 ± 6% of all germinal-layer cells expressed *emhippo* (6,491 cells analyzed), while 12 ± 3% incorporated EdU during a 5 h labeling period. Among 763 EdU+ cells analyzed, 14 ± 10% also displayed an *emhippo* hybridization signal (Fig. 3C). Thus, *emhippo* expression was detected in a subset of proliferating GC, but most *emhippo*+ cells did not incorporate EdU during the labeling period, indicating that expression is not confined to cells undergoing DNA synthesis.

We also investigated *emhippo* expression in activated protoscoleces. WISH revealed transcript signals in multiple regions of the protoscolex, including areas associated with the sucker structures and extending towards posterior regions (Fig. 3D). Thus, *emhippo* is expressed in both the proliferative metacestode germinal layer and the more differentiated protoscolex stage, consistent with a potential role of Hippo signaling in multiple aspects of larval development.

### EmYorkie expression across larval stages and cell populations

To characterize the expression of the putative Hippo pathway effector EmYorkie, we analyzed transcriptomic datasets covering different developmental stages and experimental conditions of *E. multilocularis* (Tsai et al., 2013; Herz et al., 2024; Herrmann et al., 2026). *emyorkie* transcripts were detected in primary cell cultures, metacestodes, and protoscoleces, although expression levels varied considerably between developmental stages (Fig. 4B). In primary cell cultures, *emyorkie* expression progressively increased during culture development, from approximately 13 TPM at the earliest stage to 66 TPM at the latest stage examined. Expression was particularly pronounced in protoscoleces, with transcript levels of approximately 109 TPM in non-activated and 121 TPM in activated specimens. In contrast, metacestode samples generally exhibited lower expression levels. Depletion of proliferating GC by HU treatment did not result in a marked reduction in *emyorkie* transcript abundance. Similarly, β-catenin knockdown did not significantly alter overall *emyorkie* transcript levels under the conditions examined.

**Figure 4.**
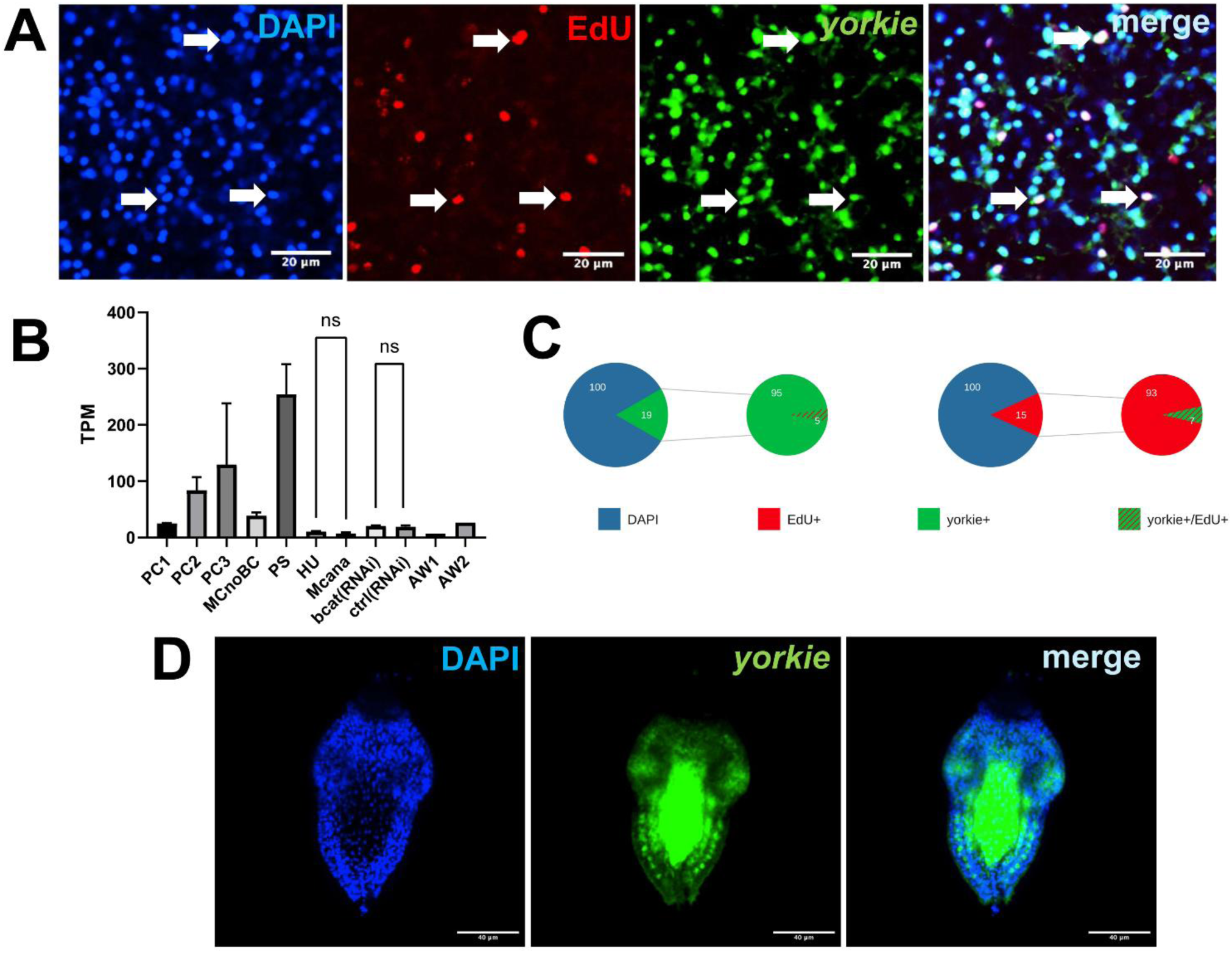
Expression of *emyorkie* in *E. multilocularis* metacestode and protoscolex. **(A)** WISH analysis of *emyorkie* expression in the germinal layer of metacestode vesicles. *emyorkie* transcripts are shown in green, EdU incorporation in red, and nuclei counterstained with DAPI in blue. Merged images illustrate the distribution of *emyorkie*-expressing cells relative to proliferating cells. Arrows indicate representative EdU-positive cells expressing *emyorkie*. Scale bars: 20 µm. **(B)** Transcriptomic expression profile of *emyorkie* across different developmental stages and experimental conditions, based on RNA-seq datasets analyzed as described by Tsai et al., 2013, Herz et al. 2024, and Herrmann et al., 2026. Expression levels are presented as transcripts per million (TPM). PC1–PC3, primary cell cultures; MCnoBC, metacestodes without brood capsules; PS, protoscoleces; HU, hydroxyurea-treated metacestodes; MCana, metacestodes maintained under anaerobic conditions; bcat(RNAi), β-catenin RNAi-treated samples; ctrl(RNAi), corresponding RNAi controls; AW1, pre-gravid adult worm; and AW2, gravid adult worm. Error bars indicate variation between samples. ns, not statistically significant. Error bars indicate variation between samples; ns, not statistically significant. **(C)** Quantification of *emyorkie* expression and EdU incorporation in the germinal layer of metacestode vesicles. The proportions of *emyorkie*-positive (19%) and EdU-positive (15%) cells were determined relative to the total number of DAPI-stained nuclei (100%). The enlarged diagrams illustrate the fractions of marker-positive cells showing overlap between *emyorkie* expression and EdU incorporation. Green, *emyorkie*-positive cells; red, EdU-positive cells; hatched areas, double-positive cells. **(D)** WISH hybridization analysis of *emyorkie* expression in activated protoscolex. *emyorkie* transcripts are shown in green and DAPI-stained nuclei in blue. The merged image illustrates the distribution of *emyorkie* expression throughout the protoscolex. Scale bars: 40 µm. All pictures represent single confocal slices.

We next investigated the spatial expression of *emyorkie* in the metacestode germinal layer by WISH, combined with a 5-hour EdU pulse to identify cells undergoing DNA synthesis. *Emyorkie*-expressing cells were distributed throughout the germinal layer without an obvious spatial clustering pattern (Fig. 4A). Quantitative analysis revealed that 19 ± 11% of all cells expressed *emyorkie* (4,941 cells analyzed), while 15 ± 5% incorporated EdU. Among 709 EdU-positive cells analyzed, only 7 ± 7% also exhibited an *emyorkie* hybridization signal (Fig. 4C). Thus, although *emyorkie* expression was detectable in a subset of proliferating GC, most transcript-positive cells were not undergoing DNA synthesis during the labeling period. Together with the transcriptomic data obtained following HU treatment, these observations indicate that *emyorkie* expression is not restricted to proliferating GC.

Finally, we examined *emyorkie* expression in activated protoscoleces. WISH revealed a regionally distributed expression pattern, with signals detected in and between the suckers and extending laterally towards posterior regions of the parasite (Fig. 4D). This distribution differed from the broadly dispersed expression observed in the metacestode germinal layer, indicating developmental stage-associated differences in the spatial distribution of *emyorkie* transcripts. Notably, in the lateral regions of activated protoscoleces, both *emhippo* and *emyorkie* displayed a distinct salt-and-pepper expression pattern, with individual transcript-positive cells interspersed among cells lacking detectable hybridization signals (Fig. 3D; Fig. 4D).

### EmHippo and EmYorkie are expressed in association with differentiated neural and muscle cells

The limited overlap between *emhippo*- or *emyorkie*-expressing cells and EdU-positive germinative cells prompted us to investigate whether these genes are also expressed in differentiated cell populations of the metacestode germinal layer. To this end, we combined WISH with immunofluorescence staining for acetylated α-tubulin, a marker of neuronal structures in *E. multilocularis* (Koziol et al., 2013), and high-molecular-weight tropomyosin, which labels muscle cells and their associated fibers (Koziol et al., 2016).

Both *emhippo*- and *emyorkie*-expressing cells were observed in close association with acetylated α-tubulin-positive neuronal structures (Fig. 5). In several instances, hybridization signals could be assigned to cells associated with neuronal processes, supporting the expression of both genes in at least a subset of differentiated neural cells. However, the extensive network of neuronal processes and the resulting diffuse immunofluorescence staining prevented reliable quantitative assessment of co-localization.

**Figure 5.**
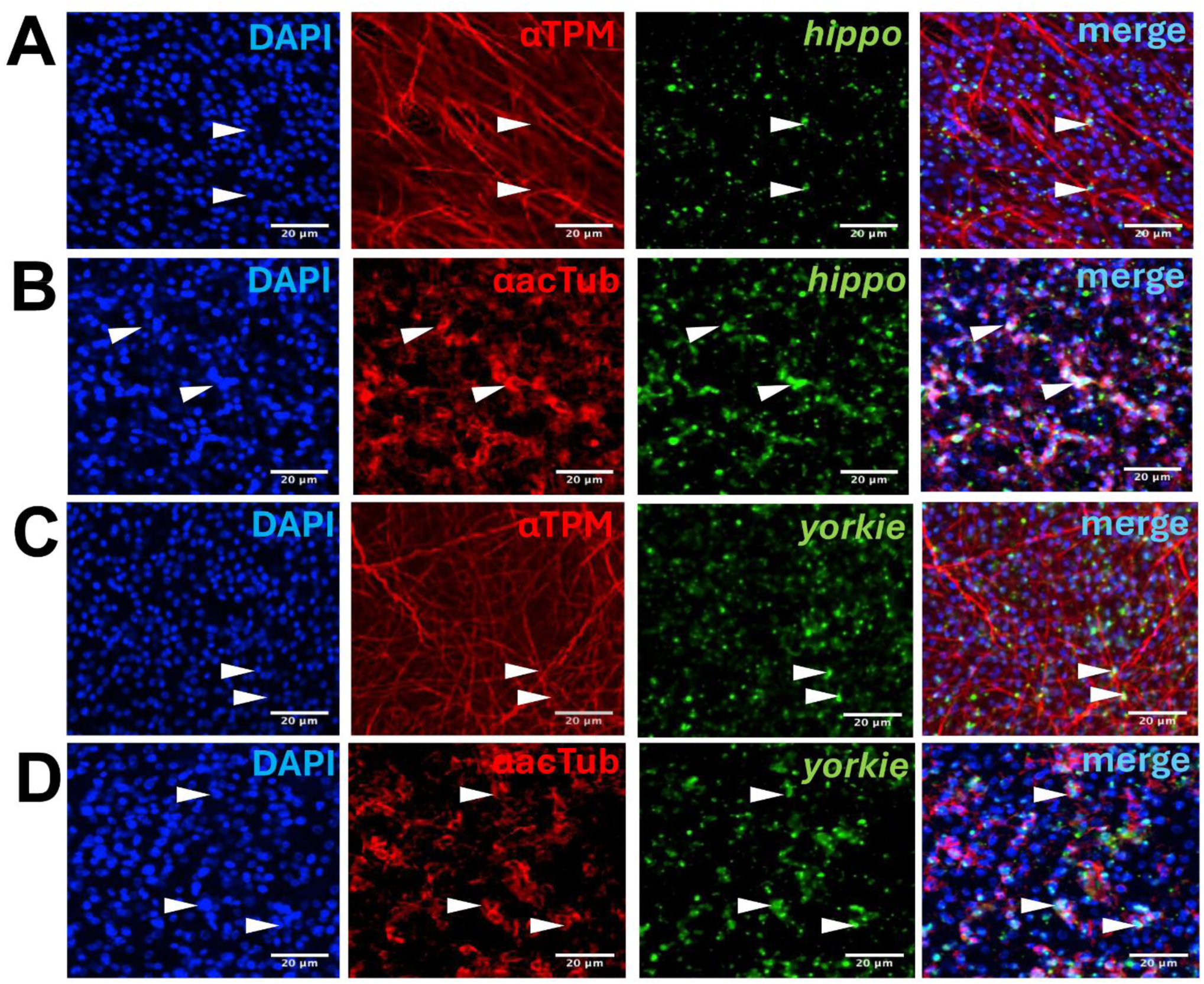
Expression of *emhippo* and *emyorkie* in muscle and neuronal cells of the *E. multilocularis* metacestode. Representative confocal images of WISH combined with immunofluorescence staining in the germinal layer of metacestode vesicles. **(A, C)** Expression of *emhippo* (A) and *emyorkie* (C) in relation to the muscle cell network. Muscle fibres were visualized using an antibody against high-molecular-weight tropomyosin (αTPM, red), while *emhippo* and *emyorkie* transcripts were detected by WISH (green). Arrowheads indicate representative transcript-positive cells associated with the muscle cell network. **(B, D)** Expression of *emhippo* (B) and *emyorkie* (D) in relation to neuronal structures. Acetylated α-tubulin immunostaining (anti-acTub, red) was used to visualize neuronal processes and associated cell bodies. Arrowheads indicate representative transcript-positive cells associated with acetylated tubulin-positive structures. In all panels, nuclei were counterstained with DAPI (blue). Individual fluorescence channels and corresponding merged images are shown. Scale bars: 20 µm in all panels. All pictures represent single confocal slices.

Similarly, *emhippo* and *emyorkie* transcripts were detected in cells associated with tropomyosin-positive muscle fibers (Fig. 5). Although the spatial separation between muscle cell bodies and their fibers complicated unequivocal assignment of individual hybridization signals, the observations were consistent with expression of both genes in a subset of differentiated muscle cells. This finding is of particular developmental interest, as muscle cells in *E. multilocularis* have been implicated in the expression of positional signaling molecules involved in larval patterning (Koziol et al., 2016; Loos et al., 2026).

Together with the EdU labeling experiments, these findings indicate that *emhippo* and *emyorkie* are expressed in multiple cellular compartments of the metacestode germinal layer, including proliferating GC and cells associated with differentiated neural and muscle structures. Thus, expression of these Hippo pathway components is not confined to the proliferative stem cell compartment.

### RNAi-mediated knockdown of EmYorkie enhances metacestode vesicle formation

To investigate the functional involvement of Hippo pathway components in *E. multilocularis* development, we performed RNA interference (RNAi) targeting *emhippo*, *emyorkie*, or both genes simultaneously in primary cell cultures. RNAi targeting *gfp* served as a negative control (Herrmann et al., 2026). Primary cell aggregates were monitored over seven days for their ability to develop metacestode vesicles and for potential changes in cell proliferation.

Knockdown efficiency was assessed by RT-qPCR at days 2 and 7 (Supplementary Fig. S6). At day 2, transcript levels of both *emhippo* and *emyorkie* were reduced following the respective RNAi treatments, including combined knockdown. At day 7, *emhippo* transcript levels remained reduced, whereas *emyorkie* expression showed partial recovery towards control levels. These measurements indicate that the extent and persistence of transcript reduction differed between the two target genes.

RNAi-mediated knockdown of *emhippo* alone did not significantly affect metacestode vesicle formation compared with the *gfp* control (Fig. 6B, C). In contrast, knockdown of *emyorkie* resulted in a pronounced increase in vesicle formation. After seven days, approximately three times as many vesicles were observed following *emyorkie* knockdown compared with control cultures. A comparable increase was detected following simultaneous knockdown of *emhippo* and *emyorkie*, indicating that additional depletion of EmHippo did not noticeably modify the phenotype associated with reduced *emyorkie* expression. The increase in vesicle formation was statistically significant compared with the control (p = 0.0088; Fig. 6B). Representative microscopy images illustrated the enhanced development of metacestode vesicles at the periphery of primary cell aggregates following *emyorkie* knockdown (Fig. 6C).

**Figure 6.**
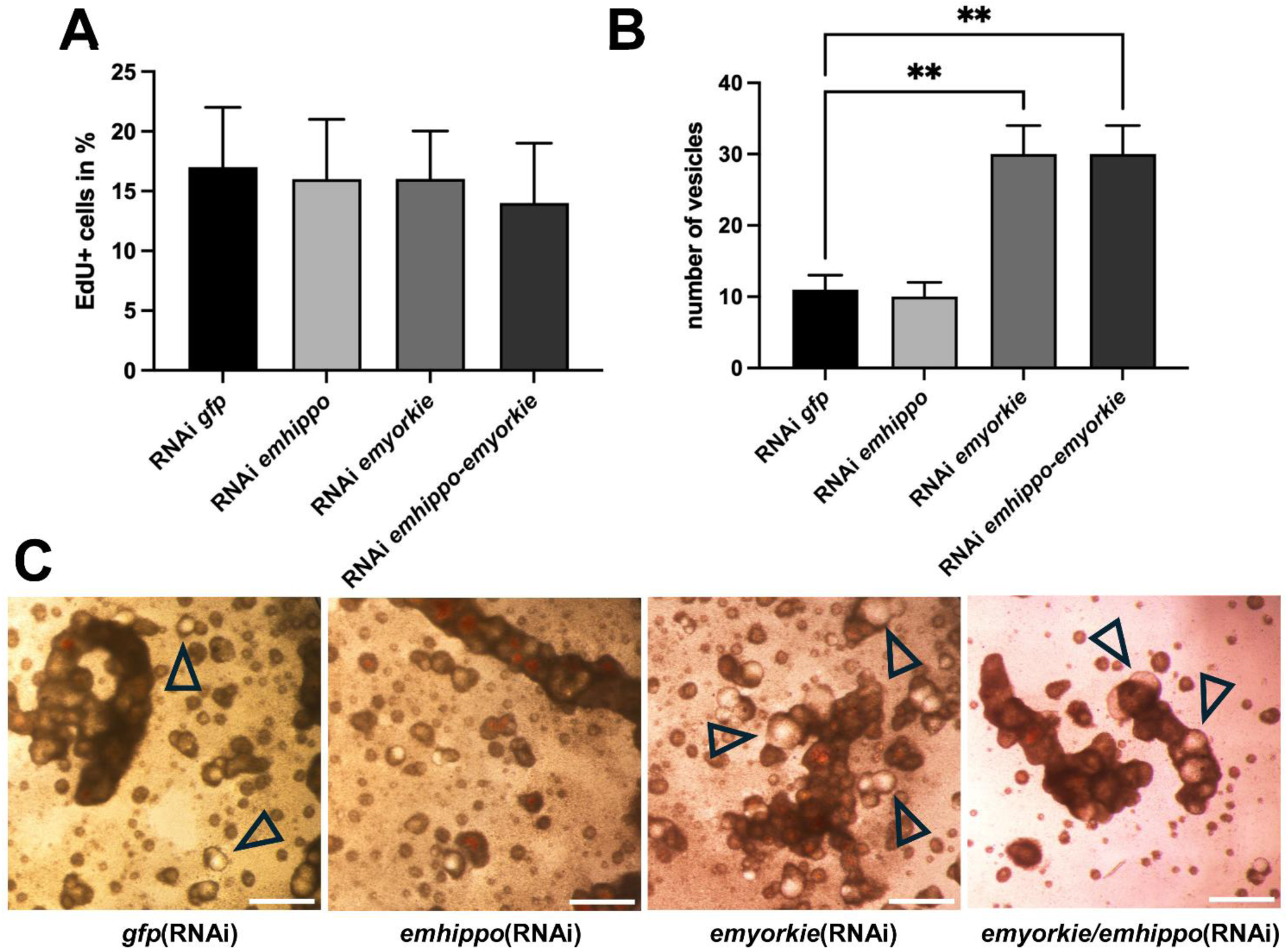
*emyorkie* RNAi enhances vesicle formation without significantly affecting germinative cell proliferation in primary cell cultures. (A) Quantification of germinative cell proliferation by EdU incorporation in primary cell cultures after 7 days of RNAi treatment targeting *emhippo*, *emyorkie*, or both genes simultaneously. RNAi targeting *gfp* served as a negative control. The proportion of EdU-positive cells was determined relative to the total cell population. No statistically significant differences were detected between treatment groups. (B) Quantification of metacestode vesicles formed from primary cell aggregates after 7 days of RNAi treatment. RNAi targeting *emyorkie*, either alone or in combination with *emhippo*, resulted in significantly increased vesicle formation compared with the *gfp*-RNAi control, whereas *emhippo* RNAi alone had no detectable effect. For (A) and (B), data are presented as mean ± SD from two independent biological experiments, each comprising three technical replicates. Technical replicates were averaged within each biological experiment before statistical analysis. Statistical significance was assessed by ANOVA followed by Dunnett’s multiple-comparisons test (**p = 0.0088; comparisons with the *gfp*-RNAi control). (C) Representative light microscopy images of primary cell aggregates following RNAi treatment targeting *gfp*, *emhippo*, *emyorkie*, or both genes simultaneously. Developing metacestode vesicles are visible at the periphery of the aggregates (open triangles), particularly following *emyorkie* RNAi. Scale bars: 200 µm.

Since Hippo signaling is frequently associated with the regulation of cell proliferation, we next investigated whether the increased vesicle formation was accompanied by changes in DNA synthesis. EdU incorporation assays performed after seven days of RNAi treatment revealed no statistically significant differences between the experimental groups (Fig. 6A). The proportion of EdU+ cells remained approximately 17% in control cultures, 16% following individual knockdown of either *emhippo* or *emyorkie*, and 14% following simultaneous knockdown of both genes.

Together, these findings indicate that RNAi targeting *emyorkie* enhances metacestode vesicle formation without a detectable increase in the proportion of EdU-positive cells at day 7. The absence of a comparable phenotype following *emhippo* RNAi, and the persistence of enhanced vesicle formation after combined knockdown, suggest that EmYorkie influences early metacestode development through processes not directly reflected in overall DNA synthesis at the time point examined. However, transient changes in proliferation at earlier stages cannot be excluded.

### XMU-MP-1 induces giant cell formation and compromises metacestode integrity

To further investigate the potential involvement of Hippo signaling in *E. multilocularis*, we employed XMU-MP-1, a pharmacological inhibitor originally developed against mammalian MST1/2 kinases (Fan et al., 2016; Triastuti et al., 2019). Sequence comparisons revealed conservation of several amino acid residues implicated in XMU-MP-1 binding within the ATP-binding pocket of mammalian MST kinases (Supplementary Fig. S1; Supplementary Fig. S9). These findings identify EmHippo as a plausible candidate target of XMU-MP-1 in *E. multilocularis*, although off-target effects on additional kinases remain possible.

Treatment of metacestode vesicles with XMU-MP-1 resulted in concentration-dependent alterations in vesicle integrity (Fig. 7A). While vesicles exposed to 1 µM largely retained their structural integrity, treatment with 10 µM caused progressive vesicle collapse. Approximately 50% of vesicles had lost structural integrity after seven days, increasing to approximately 97% after 14 days. Notably, germinal-layer tissue remained microscopically detectable even in collapsed vesicles.

**Figure 7.**
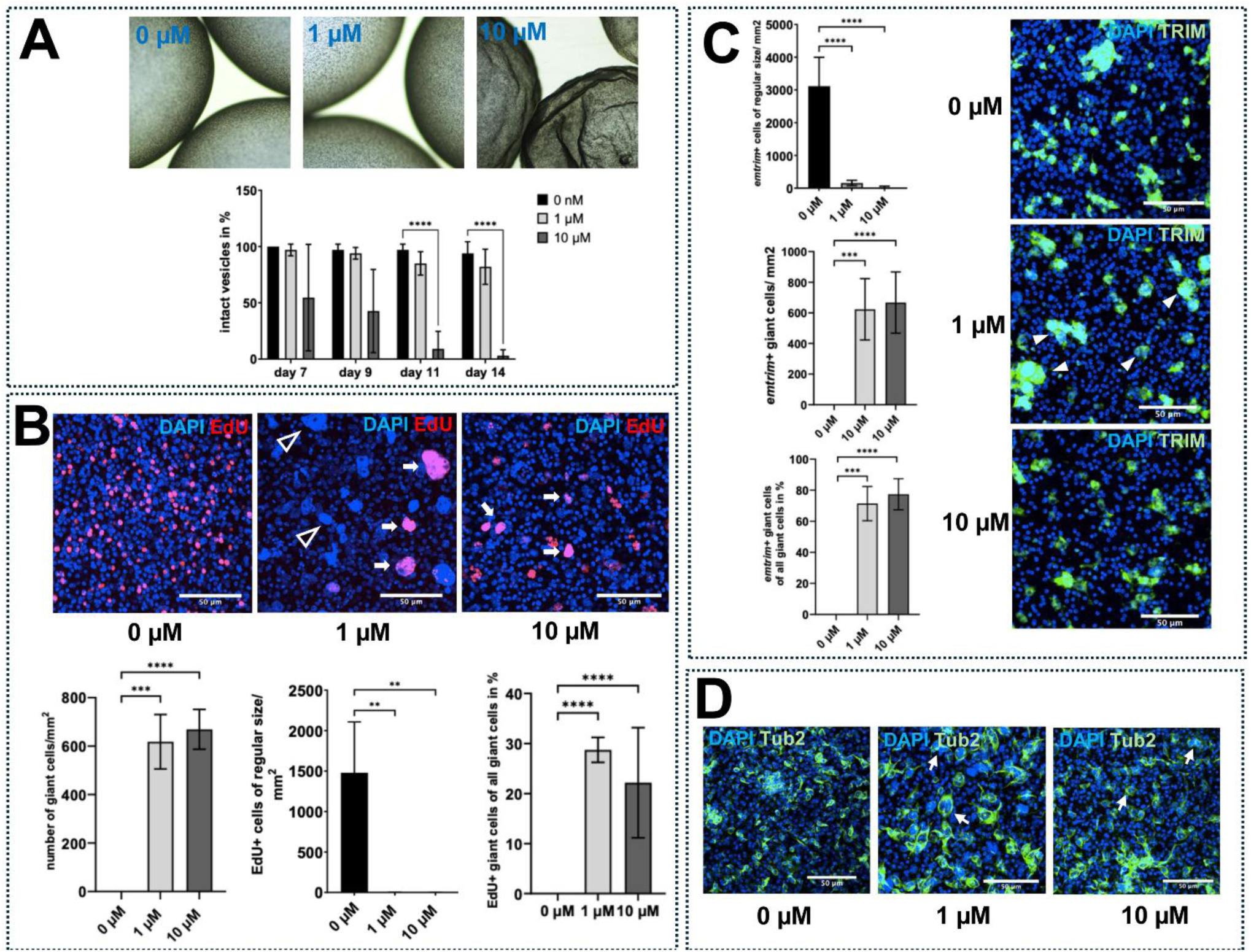
XMU-MP-1 induces giant cell formation and compromises vesicle integrity in *E. multilocularis* metacestode vesicles. (A) Effects on vesicle integrity. Upper panel: Representative brightfield images of metacestode vesicles treated with DMSO (control), 1 µM, or 10 µM XMU-MP-1 for 7 days. Treatment resulted in morphological alterations and progressive loss of vesicle integrity. Scale bars: 200 µm. Lower panel: Quantification of structurally intact metacestode vesicles following XMU-MP-1 treatment. Data represent mean ± SD from three independent experiments. Statistical analysis was performed using two-way ANOVA followed by Dunnett’s post hoc test. **(B) Giant cell formation and EdU incorporation.** Upper panel: Representative single confocal optical sections showing EdU incorporation (red) and DAPI-stained nuclei (blue) in metacestode vesicles treated with DMSO (0 µM), 1 µM, or 10 µM XMU-MP-1. EdU+ giant cells are indicated by white arrows and EdU-giant cells by open arrowheads. Scale bars: 50 µm. Lower panel: Quantification of giant cells per mm² in the germinal layer of metacestode vesicles. Data represent mean ± SD from three independent experiments. Statistical analysis was performed using one-way ANOVA followed by Dunnett’s post hoc test; ***p = 0.0001, ****p < 0.0001 compared with the DMSO control. **(C) Expression of the stem cell marker *emtrim* in XMU-MP-1-induced giant cells.** Left panel: Quantification of the proportion of *emtrim+* giant cells relative to the total number of giant cells following treatment with 1 µM or 10 µM XMU-MP-1. More than 70% of giant cells expressed *emtrim* under both treatment conditions. Data represent mean ± SD from three independent experiments. Right panel: Representative single confocal optical sections showing *emtrim*-specific WISH (green) and DAPI-stained nuclei (blue) in metacestode vesicles treated with DMSO (0 µM), 1 µM, or 10 µM XMU-MP-1 for 7 days. Scale bars: 50 µm. **(D) Detection of Tub-2 in XMU-MP-1-induced giant cells.** Representative single confocal optical sections showing Tub-2 immunostaining in metacestode vesicles treated with DMSO (0 µM), 1 µM, or 10 µM XMU-MP-1. Tub-2 was detected using a specific anti-Tub-2 antibody (green), and nuclei were counterstained with DAPI (blue). The staining reveals both enlarged cells containing a single nucleus (hypertrophic giant cells) and multinucleated giant cells. Scale bars: 100 µm. All fluorescence images shown in (B–D) represent single confocal optical sections rather than maximum-intensity projections.

In addition to these morphological alterations, XMU-MP-1 induced the formation of conspicuously enlarged cells within the germinal layer (Fig. 7B). These giant cells were already abundant following treatment with 1 µM XMU-MP-1, and their frequency exceeded 600 cells per mm² at both 1 and 10 µM. The effect was highly significant compared with DMSO-treated controls (p ≤ 0.0001). Interestingly, giant cells observed at 1 µM tended to be larger than those developing at 10 µM. Notably, pronounced giant cell formation occurred at a concentration at which metacestode vesicles largely retained their structural integrity, indicating that the cellular phenotype was not simply a consequence of generalized tissue damage. EdU incorporation demonstrated that at least a subset of these enlarged cells retained the capacity for DNA synthesis, indicating that giant cell formation was not invariably associated with complete cessation of DNA replication.

To investigate the cellular identity of the XMU-MP-1-induced giant cells, we performed WISH using *emtrim*, a marker of the germinative cell population (Koziol et al., 2015). More than 70% of giant cells expressed *emtrim* following treatment with either 1 or 10 µM XMU-MP-1 (Fig. 7C). This finding strongly supports the conclusion that the giant cells predominantly originate from the GC compartment.

We further examined the morphology of giant cells using immunostaining for the GC-associated β-tubulin isoform Tub-2 (Bergmann et al., 2026; Fig. 7D). These analyses revealed considerable heterogeneity within the giant cell population. While some enlarged cells contained a single nucleus, others displayed multiple nuclei within an enlarged cellular structure. Thus, XMU-MP-1 induced both hypertrophic and multinucleated giant cell morphologies, consistent with disturbances in normal cell-cycle progression and/or cytokinesis.

We further investigated the effects of XMU-MP-1 in primary cell cultures, which are predominantly composed of GC (Supplementary Fig. S7). Following 14 days of treatment with 1 or 10 µM XMU-MP-1, cell aggregates contained conspicuously enlarged cells, particularly at the higher concentration. At the same time, EdU incorporation was strongly reduced, indicating impaired germinative cell proliferation. Most enlarged cells showed little or no detectable EdU incorporation.

We also examined the effects of XMU-MP-1 on activated protoscoleces (Supplementary Fig. S8). Following 14 days of treatment, protoscoleces largely retained their overall morphology at both concentrations. However, treatment with 10 µM XMU-MP-1 induced enlarged cells distributed throughout the protoscolex tissue, whereas such alterations were not observed at 1 µM. Some of the enlarged cells retained the capacity to incorporate EdU, although reliable quantification was not possible.

### XMU-MP-1 disrupts mitotic progression and induces a phenotype distinct from Aurora kinase inhibition

The formation of enlarged and multinucleated germinative cells following XMU-MP-1 treatment suggested that the compound interferes with normal cell-cycle progression, potentially affecting mitosis or cytokinesis. Although XMU-MP-1 is an established inhibitor of mammalian MST1/2 kinases used to investigate Hippo signaling (Fan et al., 2016; Triastuti et al., 2019), biochemical profiling has also revealed inhibitory activity against Aurora kinases (Fan et al., 2016). This is particularly relevant because pharmacological inhibition of Aurora kinases in *E. multilocularis* has previously been shown to induce mitotic abnormalities, multinucleation, and cytokinesis defects (Cheng et al., 2019). However, sequence comparisons revealed that all eight examined residues corresponding to the reported XMU-MP-1 interaction region of mammalian MST2 are conserved in EmHippo, whereas two substitutions occur in each of the *E. multilocularis* Aurora kinases (Supplementary Fig. S9). To further characterize the pharmacological phenotype, we compared XMU-MP-1 with the Aurora A inhibitor MLN8237 at concentrations of 1 and 5 µM, corresponding to those previously used to investigate Aurora kinase inhibition in *E. multilocularis* (Cheng et al., 2019).

Both compounds reduced the number of phospho-histone H3-positive (pH3+) cells in the metacestode germinal layer, but their effects differed markedly in magnitude and concentration dependence (Fig. 8A, B). After 24 hours, treatment with 1 µM MLN8237 reduced the number of pH3+ cells from approximately 93 to 70 cells per mm², whereas treatment with 5 µM eliminated detectable pH3 staining. In contrast, exposure to only 1 µM XMU-MP-1 resulted in a complete absence of detectable pH3+ cells, compared with approximately 104 cells per mm² in the corresponding controls. This difference persisted after seven days of treatment. While MLN8237 caused a concentration-dependent reduction in pH3+ cells, no pH3+ cells were detected following XMU-MP-1 exposure at either concentration examined.

**Figure 8.**
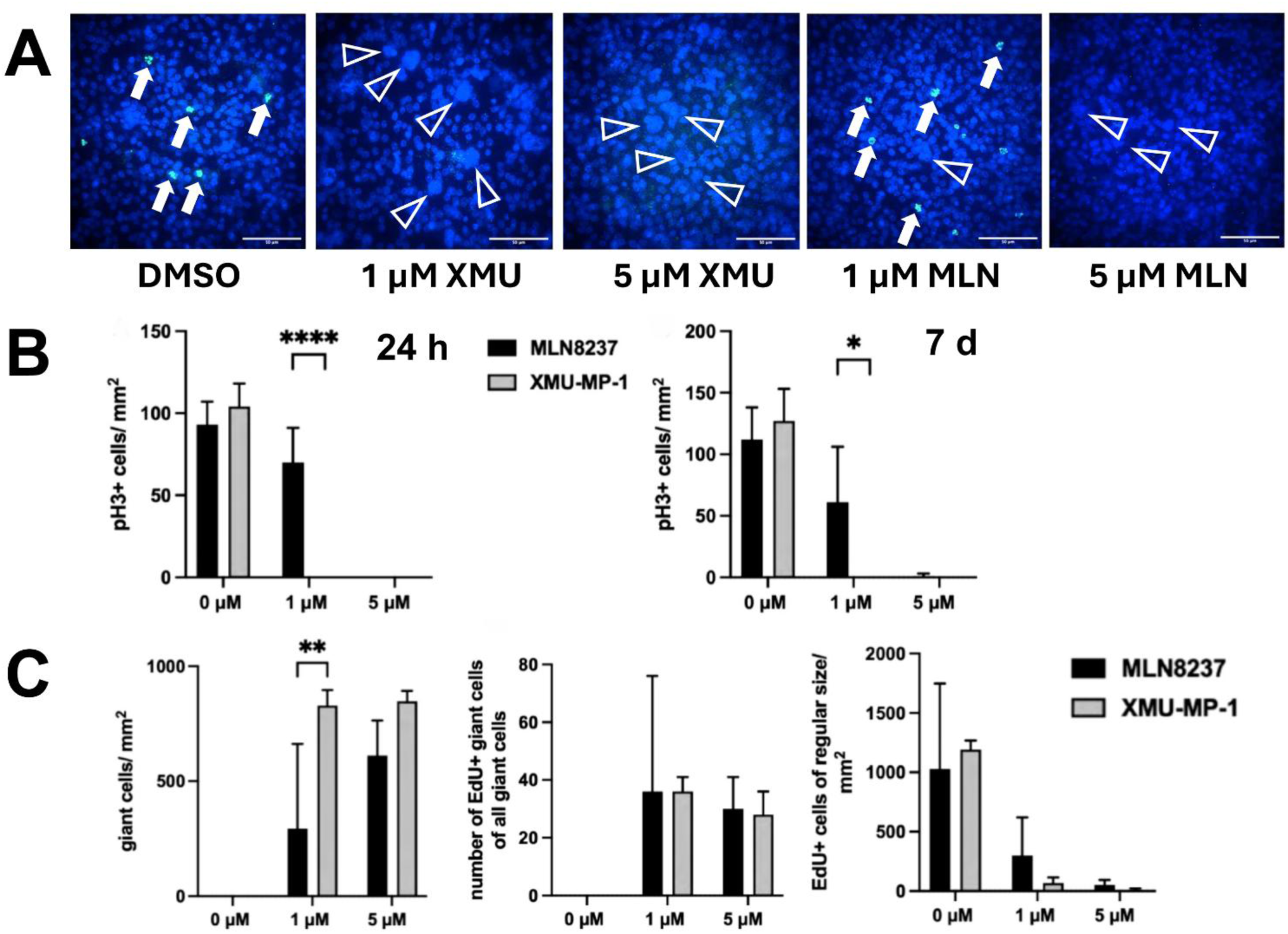
XMU-MP-1 and the Aurora kinase inhibitor MLN8237 differentially affect mitotic activity and giant cell formation in the *E. multilocularis* metacestode. **(A)** Representative single confocal optical sections showing phosphorylated histone H3 (pH3; green), a marker of mitotic cells, and DAPI-stained nuclei (blue) in metacestode vesicles treated with DMSO (control), 1 µM or 5 µM XMU-MP-1, or 1 µM or 5 µM MLN8237 for 7 days. White arrows indicate pH3+ cells, open triangles indicate giant cells. Scale bars: 50 µm. **(B)** Quantification of pH3+ mitotic cells per mm² of germinal layer following 24 h (left) or 7 d (right) of treatment with XMU-MP-1 or the Aurora kinase inhibitor MLN8237. Both inhibitors reduced mitotic activity, with XMU-MP-1 causing a more pronounced reduction at the concentrations tested. Data represent mean ± SD from three independent experiments. Statistical analysis was performed using two-way ANOVA followed by Šídák’s post hoc test; significance levels are indicated in the graph. **(C)** Quantification of giant cell formation and EdU incorporation following 7 days of treatment with XMU-MP-1 or MLN8237. Left: Number of giant cells per mm² of germinal layer. Middle: Ratio of EdU+ giant cells among all giant cells. Right: Number of regularly sized EdU+ cells per mm² of germinal layer. XMU-MP-1 induced substantially more giant cells than MLN8237, whereas giant cells generated by either treatment retained the ability to incorporate EdU. Data represent mean ± SD from three independent experiments. Statistical analysis was performed using two-way ANOVA followed by Šídák’s post hoc test; significance levels are indicated in the graphs. All fluorescence images shown in (A) represent single confocal optical sections rather than maximum-intensity projections.

We next compared giant cell formation and the capacity of enlarged cells for DNA synthesis following treatment with either compound (Fig. 8C). Both inhibitors induced enlarged or multinucleated cells, but XMU-MP-1 produced a substantially more pronounced phenotype. At 1 µM, approximately 829 giant cells per mm² were observed following XMU-MP-1 treatment, compared with 293 cells per mm² following MLN8237 exposure. Increasing the concentration to 5 µM resulted in approximately 847 giant cells per mm² following XMU-MP-1 treatment, whereas MLN8237 induced a concentration-dependent increase to approximately 611 cells per mm².

Despite these pronounced morphological abnormalities, a substantial proportion of giant cells retained the capacity to incorporate EdU. Following treatment with 1 µM of either compound, approximately 36% of giant cells were EdU+. At 5 µM, the corresponding proportions were approximately 28% for XMU-MP-1 and 30% for MLN8237. In contrast, the number of EdU+ cells with regular morphology was strongly reduced by both compounds, with XMU-MP-1 producing a particularly pronounced decrease.

Together, these findings demonstrate that XMU-MP-1 induces pronounced alterations in GC morphology and cell-cycle-associated markers, including the accumulation of enlarged cells that retain the capacity for DNA synthesis despite the absence of detectable pH3 staining. MLN8237 also induced giant-cell formation and reduced pH3 staining, but the magnitude and concentration dependence of these effects differed between the two compounds. Notably, XMU-MP-1 abolished detectable pH3 staining already at 1 µM and induced substantially more giant cells than MLN8237 at this concentration. These observations indicate that XMU-MP-1 and MLN8237 do not produce identical cellular responses under the conditions examined.

## Discussion

The Hippo signaling pathway is a central regulator of tissue growth, cell proliferation, differentiation, and organ size in metazoans (Pan, 2007; Harvey and Tapon, 2007; Pan, 2010). Its potential function in *E. multilocularis* is particularly intriguing because growth regulation differs fundamentally between developmental stages. Whereas the oncosphere, protoscolex, and adult worm develop defined body structures and dimensions, the metacestode can expand extensively through sustained growth and asexual proliferation (Koziol et al., 2014; Brehm and Koziol, 2017). In the present study, we identified the principal components of the canonical Hippo pathway in *E. multilocularis* and provide functional evidence that its downstream effector EmYorkie regulates metacestode vesicle formation. Furthermore, pharmacological interference with XMU-MP-1 induced profound alterations in GC morphology and cell-cycle regulation. Together, these findings implicate Hippo-associated signaling in the coordination of growth and developmental processes in the larval stages of *E. multilocularis*.

The identification of EmHippo, EmSalvador, EmMats, EmWarts, EmYorkie, and EmScalloped is consistent with the ancient evolutionary origin of the Hippo pathway (Hilman and Gat, 2011; Phillips et al., 2024). EmYorkie was particularly difficult to identify because of its pronounced sequence divergence, yet it retains a WW domain and candidate Warts/LATS phosphorylation motifs. Importantly, the reciprocal yeast two-hybrid interaction between EmYorkie and EmScalloped provides experimental support for a conserved transcriptional regulatory module. In other metazoans, Yorkie/YAP proteins interact with Scalloped/TEAD transcription factors to regulate the expression of genes involved in proliferation, survival, and differentiation (Wu et al., 2008; Zhao et al., 2008; Misra and Irvine, 2018). The conservation of this interaction in *E. multilocularis*, despite considerable divergence of the EmYorkie sequence, strongly supports its functional identity as a Yorkie ortholog. Whether EmWarts directly phosphorylates EmYorkie at the predicted regulatory sites, and which downstream genes are controlled by the EmYorkie–EmScalloped complex, remain important questions for future investigations.

Our expression analyses suggest that Hippo signaling may function in several cellular and developmental contexts. Both *emhippo* and *emyorkie* transcripts were detected in EdU+ and EdU-cells and in association with neural and muscle structures. Such a distribution is compatible with the broad roles of Hippo pathway components in other metazoans, where their functions extend beyond the regulation of stem cell proliferation (Piccolo et al., 2014; Misra and Irvine, 2018). Particularly noteworthy was the substantially higher abundance of *emyorkie* transcripts in protoscoleces compared with metacestode vesicles. This difference may reflect distinct requirements for growth regulation and tissue organization in these developmental stages. In addition, the association of *emhippo* and *emyorkie* expression with muscle structures may be relevant to the regulation of developmental patterning. In planarians, muscle cells provide positional information through the expression of positional control genes, including components of the WNT signaling system (Witchley et al., 2013). Comparable expression of developmental regulators in muscle cells has been described in *Echinococcus* (Koziol et al., 2014; Loos et al., 2026). Thus, Hippo signaling may participate not only in the regulation of individual cell behavior but also in the coordination of tissue organization and developmental signals.

The most prominent genetic phenotype was observed following *emyorkie* RNAi, which significantly increased vesicle formation in primary cell cultures without significantly affecting the proportion of EdU+ cells. This suggests that EmYorkie depletion may promote the differentiation of existing GC towards metacestode vesicle formation rather than stimulating their proliferation. Interestingly, inhibition of WNT/β-catenin signaling has previously been shown to promote anteriorization in *E. multilocularis* (Herrmann et al., 2026), whereas the increased vesicle formation following *emyorkie* knockdown might indicate a shift towards posterior developmental identity. Functional interactions between Hippo/YAP and WNT/β-catenin signaling have been demonstrated in other metazoans (Azzolin et al., 2014), and studies in planarians have implicated Hippo pathway components in tissue patterning and positional information (Lin and Pearson, 2014; de Sousa et al., 2018; Doddihal et al., 2024). Together, these observations suggest that Hippo/Yorkie and WNT signaling may interact in regulating developmental fate decisions in *E. multilocularis*. Although *emhippo* and *emyorkie* expression remained unchanged following β-catenin knockdown (Herrmann et al., 2026), interactions at the level of protein activity or downstream signaling remain possible. Combined perturbation of both pathways could help clarify their respective contributions to anterior–posterior patterning and metacestode development.

In contrast to *emyorkie* knockdown, RNAi targeting *emhippo* did not produce a detectable developmental phenotype, despite a reduction in transcript abundance. This finding should not necessarily be interpreted as evidence that EmHippo is dispensable for metacestode development. RNAi in *E. multilocularis* is technically much more challenging than in planarians, where prolonged and repeated dsRNA administration can produce sustained depletion of target transcripts. In our experiments, knockdown was transient and target transcript levels partially recovered during the observation period. Moreover, reduction of mRNA abundance does not necessarily result in a corresponding reduction in protein activity, particularly when the targeted protein is relatively stable.

The stability of EmHippo has not yet been determined, whereas Yorkie/YAP proteins are known to undergo phosphorylation-dependent regulation and proteasomal degradation (Zhao et al., 2010). Differences in protein turnover could therefore contribute to the distinct RNAi phenotypes.

Pharmacological treatment with XMU-MP-1 revealed a second, particularly striking phenotype. The compound induced numerous enlarged cells within the germinal layer, even at concentrations that largely preserved metacestode vesicle integrity. The vast majority of these giant cells expressed the GC marker *emtrim* (Koziol et al., 2015) and immunostaining for the GC-associated β-tubulin isoform Tub-2 (Bergmann et al., 2026) revealed both enlarged mononucleated and multinucleated cells. These observations indicate that the giant cells originate from the GC compartment rather than representing nonspecific enlargement of differentiated cells. Their appearance before extensive vesicle collapse further indicates that giant cell formation is not merely a secondary consequence of generalized tissue damage. At higher concentrations and following prolonged exposure, XMU-MP-1 caused progressive loss of vesicle integrity, consistent with the importance of continuous GC activity for metacestode tissue maintenance (Schubert et al., 2014).

The combination of continued DNA synthesis, the absence of detectable pH3+ cells, and the formation of enlarged GC suggests that XMU-MP-1 profoundly disrupts normal cell-cycle progression. Interestingly, the giant cell population comprised both mononucleated and multinucleated cells, indicating that individual germinative cells may respond differently to inhibitor treatment. One possible explanation is that the cellular outcome depends on the cell-cycle phase at the time of exposure. Given the relatively long cell cycle of *E. multilocularis* GC, estimated at approximately three to four days (Cheng et al., 2017; Herz et al., 2024), cells entering treatment at different stages might undergo distinct alterations in cell growth, DNA replication, and nuclear division. Thus, the heterogeneous giant cell phenotype may reflect different consequences of a common disturbance in cell-cycle regulation.

Such an interpretation is biologically plausible in view of the established involvement of Hippo pathway components in cell-cycle regulation and the formation of enlarged or polyploid cells in other organisms. For example, YAP depletion has been associated with abnormalities in spindle organization and cytokinesis in mammalian cells, resulting in multinucleation and polyploidy (Bui et al., 2016). Furthermore, Hippo pathway components have been implicated in the differentiation of trophoblast stem cells into polyploid giant cells, a process involving endoreduplication (Basak and Ain, 2022). These observations do not establish that identical mechanisms operate in *E. multilocularis* but demonstrate that interference with Hippo-associated signaling can produce cellular abnormalities resembling those observed in our experiments.

An important consideration in interpreting the XMU-MP-1 phenotype is the pharmacological specificity of the compound. XMU-MP-1 is a well-established ATP-competitive inhibitor of mammalian MST1/2 kinases and has been widely employed to investigate Hippo pathway function in cellular and animal models (Fan et al., 2016; Triastuti et al., 2019). However, biochemical profiling has also demonstrated inhibitory activity against additional kinases, including Aurora kinases (Fan et al., 2016). This is particularly relevant because inhibition of Aurora kinases in *E. multilocularis* has previously been shown to induce mitotic abnormalities, multinucleation, and cytokinesis defects (Cheng et al., 2019). The occurrence of partially overlapping cellular phenotypes following Hippo-associated and Aurora kinase inhibition is therefore not unexpected, since both kinase systems participate in interconnected mechanisms regulating cell growth, mitosis, and cytokinesis. Nevertheless, the comparison with the Aurora kinase inhibitor MLN8237 revealed important differences.

Cheng et al. (2019) reported a significant increase in the proportion of pH3+ GC following treatment of *E. multilocularis* metacestode vesicles with 1 µM MLN8237 for 24 h, consistent with an accumulation of cells displaying a mitotic marker after preferential inhibition of Aurora A. Under our experimental conditions, we did not observe a corresponding increase. Instead, the mean number of pH3+ cells per mm² was numerically lower following treatment with 1 µM MLN8237 at both 24 h and 7 days, although these differences were not statistically significant compared with the respective controls. Importantly, the two studies employed different quantification approaches: Cheng et al. (2019) determined the proportion of pH3+ cells, whereas we measured their density per unit area of germinal layer. Differences in experimental conditions may also have contributed to the divergent observations. In contrast to MLN8237, treatment with 1 µM XMU-MP-1 resulted in a complete absence of detectable pH3+ at both time points. Thus, under the conditions examined here, preferential Aurora A inhibition did not reproduce the pronounced loss of pH3 staining observed following XMU-MP-1 treatment.

The interpretation of these findings requires consideration of the molecular basis of histone H3 phosphorylation. Phosphorylation of histone H3 at serine 10 during mitosis is predominantly associated with Aurora B activity rather than Aurora A. Cheng et al. (2019) showed that higher concentrations of MLN8237 can also interfere with Aurora B-associated histone H3 phosphorylation. Consequently, the absence of detectable pH3 staining following XMU-MP-1 treatment does not necessarily indicate a complete absence of mitotic activity. It could reflect interference with mitotic progression, reduced histone H3 phosphorylation, or a combination of these effects. Our experiments do not directly distinguish between these possibilities. Nevertheless, the marked differences in pH3 staining and giant-cell formation between XMU-MP-1 and MLN8237 indicate that the two compounds elicit distinct cellular responses under the conditions examined. Although Aurora kinase inhibition may contribute to the XMU-MP-1 phenotype, its effects cannot be fully explained by assuming a cellular response identical to that induced by MLN8237.

Several observations make EmHippo a plausible candidate target of XMU-MP-1 in *E. multilocularis*. First, XMU-MP-1 was developed as an inhibitor of mammalian MST1/2 kinases (Fan et al., 2016), providing a pharmacological rationale for investigating its effects on the corresponding parasite kinase. Second, all eight examined residues corresponding to the reported XMU-MP-1 interaction region of mammalian MST2 are conserved in EmHippo, whereas two substitutions occur in each of the *E. multilocularis* Aurora kinases. Although these sequence features are consistent with the possibility of XMU-MP-1 binding to EmHippo, they do not establish inhibitor affinity or selectivity. Third, XMU-MP-1 produced cellular effects that differed from those of MLN8237, particularly the pronounced loss of detectable pH3 staining and the greater induction of giant cells at low concentrations. Moreover, giant-cell formation was observed under conditions in which metacestode vesicles initially retained their structural integrity, making generalized tissue damage an unlikely sole explanation for the phenotype. Taken together, these findings are compatible with a contribution of EmHippo inhibition to the observed effects. However, the available data do not establish EmHippo as the primary molecular target of XMU-MP-1, and contributions from Aurora kinases or other targets remain possible.

Direct biochemical validation of EmHippo inhibition would provide an important further test of this interpretation. However, establishing a physiologically informative EmHippo kinase assay is technically challenging because it would ideally require a suitable recombinant substrate, such as EmWarts, and potentially regulatory components such as EmSalvador. Attempts to monitor EmWarts phosphorylation using phospho-specific antibodies did not yield a detectable signal in our hands, potentially reflecting insufficient conservation of the corresponding phosphorylation-site epitopes. This negative result therefore cannot be interpreted as evidence for absent EmHippo activity. Future biochemical studies, phosphoproteomic approaches, or genetic experiments enabling more sustained depletion of Hippo pathway components will be necessary to define the direct molecular targets of XMU-MP-1 and establish the downstream signaling events responsible for giant cell formation.

The effects of XMU-MP-1 differed considerably between developmental systems. In metacestode vesicles, treatment induced numerous enlarged GC, a proportion of which retained the capacity for DNA synthesis. In contrast, primary cell cultures, which are predominantly composed of GC (Koziol et al., 2014), showed a pronounced reduction in EdU incorporation, whereas conspicuous cellular enlargement in protoscoleces was mainly observed at higher inhibitor concentrations. The different responses of GC in metacestode tissue and primary cell cultures suggest that the cellular consequences of inhibitor treatment may depend not only on cell identity but also on developmental state and tissue context. Differences in compound accessibility or sensitivity may also contribute. The distinct expression patterns of *emyorkie* in metacestodes and protoscoleces are consistent with the possibility that Hippo-associated signaling fulfils different functions during larval development. However, the molecular targets of XMU-MP-1 and their relative contributions to the observed phenotypes remain to be established in each developmental system.

In conclusion, our findings identify conserved components of the Hippo signaling machinery in *E. multilocularis* and provide evidence for a role of EmYorkie in early larval development. The enhanced formation of metacestode vesicles following *emyorkie* knockdown, without a detectable increase in EdU incorporation at the time point examined, suggests that EmYorkie influences developmental processes not directly reflected in overall DNA synthesis. In parallel, pharmacological treatment with XMU-MP-1 revealed pronounced alterations in GC morphology and cell-cycle-associated markers, including the formation of enlarged and multinucleated cells that can retain DNA synthesis activity. Although the precise molecular targets responsible for these pharmacological effects remain unresolved, the conservation of the predicted XMU-MP-1 interaction region in EmHippo provides a rationale for further investigation of this kinase. Together, the genetic and pharmacological findings highlight the potential importance of Hippo-associated signaling in larval development and establish a foundation for investigating its relationship with germinative-cell regulation in parasitic flatworms.

## Acknowledgments

The authors wish to thank Dirk Radloff for excellent technical assistance.

## Ethics Statement

This study was performed in strict accordance with German (Deutsches Tierschutzgesetz, TierSchG, version from Dec-9-2010) and European (European directive 2010/63/EU) regulation on the protection of animals. For experiments involving *Meriones unguiculatus*, the protocol was approved by the Ethics Committee of the Government of Lower Franconia under permit numbers 55.2.2-2532-2-1479-8.

## Funding

This work was supported by the Bayerische Transformations- und Forschungsstiftung (https://www.forschungsstiftung.bayern.de; grant [AZ-1341-18] (to KB) and the Manfred Wellhöfer foundation (Schwarzenbruck, Germany; grant 824000) (to KB). The authors declare that they have no competing interests.

## Data Availability

All data supporting the findings of this study are included in the manuscript and its supplementary materials. Additional data are available from the corresponding author upon reasonable request.

## Notes

### Competing Interest Statement

The authors have declared no competing interest.

