## Supplementary Figures S1 - S9 for "Inhibition of Hippo signalling disrupts germinative cell-cycle progression and larval development in *Echinococcus multilocularis*"

Figure S1

|  |  |  |
| --- | --- | --- |
| EmHippo | MAQKAGPPSEFLQSTVSNCTWDIGYLREMKKLESELSKPPSKVLEVICLKGKSGYSV | 60 |
| sp:HIPPO_DROME | ---MSEPEVTSVVDMKSPNISSSCSFFKLKKLSEESLLQPPEKVFDIMYKLGEGSYGSVY | 57 |
| sp:STK4_HUMAN | -----METVQ-----LRNPPRRQLKKLDEDSLTKQPEEVFDVLEKLGEGSYGSVY | 45 |
| sp:STK3_HUMAN | -----ME-----QPPAPKSKLKKLSEDSLTKQPEEVFDVLEKLGEGSYGSVF | 42 |
|  | : : * * . . * : * : : : : * * : * * * * : |  |
| EmHippo | KAKHKASGTVVAVKKVPVDSDLSEIVKEISIMQQCDSPYIVKCFGSLFDNQDLWICMEYC | 120 |
| sp:HIPPO_DROME | KAVHKESSSIVAIKLVPVESDLHEIIKEISIMQQCDSPYVVRYYGSYFKQYDLWICMEYC | 117 |
| sp:STK4_HUMAN | KAIHKETGQIVAIKQVPVESDLQEIIKEISIMQQCDSPHVVKYYGSYFKNTDLWIVMEYC | 105 |
| sp:STK3_HUMAN | KAIHKESGQVVAIKQVPVESDLQEIIKEISIMQQCDSPYVVKYYGSYFKNTDLWIVMEYC | 102 |
|  | ** ** : . : * : * * * : * * * * * : : * : * : * * * * * |  |
| EmHippo | GAGSVADIMRLRNQTLTEEEIGTILKYSLLGLDYLHQMRKIHRDIKAGNILLTSSGMAKL | 180 |
| sp:HIPPO_DROME | GAGSVSDIMRLRKKTLTEDEIATILSDTLQGLVYLHLRRKIHRDIKAANILLNTEGYAKL | 177 |
| sp:STK4_HUMAN | GAGSVSDIIRLRNKTLTEDEIATILQSTLKGLEYLHFMRKIHRDIKAGNILLNTEGHAKL | 165 |
| sp:STK3_HUMAN | GAGSVSDIIRLRNKTLIEDEIATILKSTLKGLEYLHFMRKIHRDIKAGNILLNTEGHAKL | 162 |
|  | * * * * : * : * * : * * : * * * * : * * * * * * * * * * : . * * * |  |
| EmHippo | ADFGVAGQLVDTLAKRNTVIGTFFWMAPEVIQEVGYNCSDIWSLGITAIEMGDGKPPGL | 240 |
| sp:HIPPO_DROME | ADFGVAGQLTDTMAKRNTVIGTPFWMAPEVIEEIGYDCVADIWSLGITALEMAEGKPPYG | 237 |
| sp:STK4_HUMAN | ADFGVAGQLTDTMAKRNTVIGTPFWMAPEVIQEIGYNCVADIWSLGITAIEMAEGKPPYA | 225 |
| sp:STK3_HUMAN | ADFGVAGQLTDTMAKRNTVIGTPFWMAPEVIQEIGYNCVADIWSLGITSIEMAEGKPPYA | 222 |
|  | * * * * * : * : * * * * * * * * * * : * : * * * * * * * * : * * * * |  |
| EmHippo | DEHPMRALFMIPSQPSPRLRDETAWSVDFVAFVNSCLARAPEARPSAAALLQTNFIRGAK | 300 |
| sp:HIPPO_DROME | EIHPMRAIFMIPQKPPPSFREPDRWSTEFIDFVSKCLVKEPDDRATATELLEHEFIRNAK | 297 |
| sp:STK4_HUMAN | DIHPMRAIFMIPTNPPPTFRKPELWSDNFTDFVKQCLVKSPEQRATATQLLQHPFVRSAK | 285 |
| sp:STK3_HUMAN | DIHPMRAIFMIPTNPPPTFRKPELWSDDFTDFVKKCLVKNPEQRATATQLLQHPFIKNAK | 282 |
|  | : * * * * : * * * : * * : * * : * : * : * : * : * : * : * * |  |
| EmHippo | SCYILLPLIEESNRLREKRLNDSKAAAASTTSPQTPFAKASVKARARPQEVIDIVEEETS | 360 |
| sp:HIPPO_DROME | HRSILKPMLEETCAIREQQRANRSFGGVLAASQAKSLAT--Q-ENGMQHITDN--AFME | 352 |
| sp:STK4_HUMAN | GVSILRDLINEAMDVKLKRQESQQ-----REVDQDDE--ENSEDEM----- | 325 |
| sp:STK3_HUMAN | PVSILRDLITEAMEIKAKRHEEQQ-----RELEE-EE-ENSEDEL----- | 321 |
|  | ** : : * : : : : . . . . . : : |  |
| EmHippo | NSGTMIQR-----SGSSSMIVKDEG---AGATGDASETLIVLNQSA-----RGGT- | 402 |
| sp:HIPPO_DROME | DPGTLVPEKFGEYQQSSASDATMIAHAEQGVDEGTLGPG--GLRNLSKAAAPAAASSAAS | 410 |
| sp:STK4_HUMAN | DSGTMVRA-----VG--DEM-----G--TVRVASTM-----TDGA- | 351 |
| sp:STK3_HUMAN | DSHTMVKT-----SV--ESV-----G--TMRATSTM-----SEGA- | 347 |
|  | : * : : : . : . : . : : |  |
| EmHippo | -----SGQDGATTMVINEDFGAEDDTECSVVIHAGDSMLEKDPGAENAGTLT | 449 |
| sp:HIPPO_DROME | PLDMPAVDSGTMVELESNLGTMVINSDSDDSTTAKN-----NDDQKPRNRYRPQFL | 461 |
| sp:STK4_HUMAN | ---NTMIEHDDT-LPSQLGTMVINAEDEE-EEGTM-----KRDETMQPAKPSFL | 396 |
| sp:STK3_HUMAN | ---QTMIEHNSTMLESDLGTMVINSEDEEEEDGTM-----KRNATSPQVQRPSFM | 394 |
|  | . * * * * : : |  |
| EmHippo | QNFARQLAIGNRCKESENVAAAVAAAQRDCGAIP-----SPTSPPSASESDSCSS | 499 |
| sp:HIPPO_DROME | EHFDRKNA-GDGRGDEKPIATEYSPAAAEQQQQQQQQQQQQQQDEQHLASGANDLNNWEH | 520 |
| sp:STK4_HUMAN | EYFEQKEK-ENQINS-----FGK | 413 |
| sp:STK3_HUMAN | DYFDKQDF-KNKSHE-----NCNQ | 412 |
|  | : * : : : . |  |
| EmHippo | SPSHHHH-----HHHLLLPHTSGQRPFAPFSRAFHE-----SGALLQQ---QQQOGMS | 544 |
| sp:HIPPO_DROME | NMEMQFQQISAINQYGLQQHQQQQQVLMAYPLMNEQLIALNNQPNLLLSNAAPMGQQGIP | 580 |
| sp:STK4_HUMAN | SVP-----GP----- | 418 |
| sp:STK3_HUMAN | NMH-----EPFP-----MSK---- | 422 |
|  | . |  |
| EmHippo | PFDGGGEVPPPCHANA-----LALANEPGGLSKLSYSELEQLLVNLGRDLETELRNLA | 597 |
| sp:HIPPO_DROME | -AA-APAQPPPAYQNQHMHTQSHAYVEGEFEFLKFLTFDDLNQRLCNIDHEMELEIEQLN | 638 |
| sp:STK4_HUMAN | -----LKNSSDWKIPQDGDYEFLKSWTVEDLQKRLLALDPMMEQEIEEIR | 463 |
| sp:STK3_HUMAN | -----NVFPDNWKVPQDGDFDFLKNLSLEELQMRLKALDPMMEREIEELR | 467 |
|  | : : * . : : * : * : * : : |  |
| EmHippo | IRYRHKRQPLLDAIAEKTAALAP----- | 620 |
| sp:HIPPO_DROME | KKYNAKRQPIVDAMNAKRKRQQNINNNLIKI | 669 |
| sp:STK4_HUMAN | QKYQSKRQPILDAIEAKKRRQQNF----- | 487 |
| sp:STK3_HUMAN | QRYTAKRQPILDAMDAKKRRQQNF----- | 491 |
|  | : * * * * : * * : * |  |

**Figure S1: Sequence features and homologies of EmHippo.** Depicted is an amino acid sequence comparison between full length EmHippo (EmuJ\_000067500) and *D. melanogaster* Hippo (UniProt Q8T0S6) as well as human STK4 (MST1; Uniprot Q13043) and STK3 (MST2; Uniprot Q13188). Sites of perfect alignment (\*) as well as groups of strong (:) or weak (.) similarity are marked below the alignment. The predicted serine/threonine kinase and MST1\_SARAH domains of EmHippo are marked in blue and violet, respectively. Blue triangles above the alignment indicate residues involved in XMU-MP-1 binding.

Figure S2

|  |  |  |
| --- | --- | --- |
| EmMATS | MKRRFWCVVNPFEVKETSRRCNRAGSRTFRQKKNSGSECNRNGGAAAIGATVSGSGGVTT | 60 |
| sp:MOB1_DROME | -----MDFLFGSRSSKTFKPKKNIP----- | 20 |
| sp:MOB1A_HUMAN | -----MSFLFSSRSSKTFKPKKNIP----- | 20 |
| sp:MOB1B_HUMAN | -----MSFLFGSRSSKTFKPKKNIP----- | 20 |
|  | . . . *: **: *** |  |
| EmMATS | NANGVCILPPEAAGTLGSGDLKQAVRLPTGEDLCEWVAVNTVEFFQQINMLYGTLAEFCT | 120 |
| sp:MOB1_DROME | EGTHQYDLMKHAAATLGSGNLRNAVALPDGEDLNEWVAVNTVDFFNQINMLYGTITEFCT | 80 |
| sp:MOB1A_HUMAN | EGSHQYELLKHAEATLGSGNLRQAVMLPEGEDLNEWIAVNTVDFFNQINMLYGTITEFCT | 80 |
| sp:MOB1B_HUMAN | EGSHQYELLKHAEATLGSGNLRMAVMLPEGEDLNEWVAVNTVDFFNQINMLYGTITDFCT | 80 |
|  | :.. * . * .*****: : ** ** ***** **:*****:***:*****: : :*** |  |
| EmMATS | DTTCPIMSAGPRYEYYWADGQTVKKPVKCSAPRYINNLMTWTQNQLENEAIFPSKIGVPF | 180 |
| sp:MOB1_DROME | EETCGIMSAGPKYEYHWADGLTVKKPIKCSAPKYIDYLMTWVQDQLDDETLFPSKIGVPF | 140 |
| sp:MOB1A_HUMAN | EASCPVMSAGPRYEYHWADGTNIKKPIKCSAPKYIDYLMTWVQDQLDDETLFPSKIGVPF | 140 |
| sp:MOB1B_HUMAN | EESCPVMSAGPKYEYHWADGTNIKKPIKCSAPKYIDYLMTWVQDQLDDETLFPSKIGVPF | 140 |
|  | : : * :*****:***:***** . :*****:*****:***: *****.*:***: : : :***** |  |
| EmMATS | PPNFLKVAKAILKRLFRVYAHIYNVHFREVQLLDEATHLNTSFKHFVYFVLEFDLVRSVE | 240 |
| sp:MOB1_DROME | PKNFHSSAKTILKRLFRVYAHIYHQHFTEVVTLGEEAHLNTSFKHFIFFVQEFNLIERRE | 200 |
| sp:MOB1A_HUMAN | PKNFMSVAKTILKRLFRVYAHIYHQHFDSVMQLQEEAHLNTSFKHFIFFVQEFNLIDRRE | 200 |
| sp:MOB1B_HUMAN | PKNFMSVAKTILKRLFRVYAHIYHQHFDPVIQLQEEAHLNTSFKHFIFFVQEFNLIDRRE | 200 |
|  | * ** . **:*****:*****: ** * * * :*****: : ** **: * : * |  |
| EmMATS | LEPLQPVIDVLTAPKTDRPTGSNNANSGASHHQSGGKNTNGVVQAEPMPPSPPVLPQPTA | 300 |
| sp:MOB1_DROME | LAPLQELIDKLTAKDERQI----- | 219 |
| sp:MOB1A_HUMAN | LAPLQELIEKLGSKDR----- | 216 |
| sp:MOB1B_HUMAN | LAPLQELIEKLTSKDR----- | 216 |
|  | * *** :*: * : . |  |
| EmMATS | SSTRM | 305 |
| sp:MOB1_DROME | ----- | 219 |
| sp:MOB1A_HUMAN | ----- | 216 |
| sp:MOB1B_HUMAN | ----- | 216 |

Figure S2: Sequence features and homologies of EmMats. Depicted is an amino acid sequence comparison between full length EmMATS (EmuJ\_000189500) and *D. melanogaster* MOB1 (UniProt Q95RA8) as well as human MOB1A (Uniprot Q9H8S9) and MOB1B (Uniprot Q7L9L4). Sites of perfect alignment (\*) as well as groups of strong (:) or weak (.) similarity are marked below the alignment. The predicted Mob1\_phocein domain of EmMATS is marked in blue.

|  |  |  |
| --- | --- | --- |
| EmWarts_CT | FIRIKRLGVGAFGWVWLVRKKEN-RQLYAMKILKKRDVVRRRQLAHVQAERDILAEADSD | 59 |
| sp:WARTS_DROME_CT | FVKLKPIGVGAFGEVTLSKIDTNSHLYAMKTLLRKADVLKRNVAVHVKAERDILAEDNND | 60 |
| sp:LATS2_HUMAN_CT | FVKIKTLGIGAFGEVCLACKVDT-HALYAMKTLRKKDVLNRNQVAHVKAERDILAEADNE | 59 |
| sp:LATS1_HUMAN_CT | FVKIKTLGIGAFGEVCLARKVDT-KALYATKTLRKKDVLLRNQVAHVKAERDILAEADNE<br>*:::*: *:**** * *. * :. . *** * *: * **: *.*:***:*****.:. | 59 |
| EmWarts_CT | WVVKLFFSFQDSHALYLVMEYIPGGDMMSLLIKKEIFEEPLARFYTAELTLALGSVHELG | 119 |
| sp:WARTS_DROME_CT | WVVKLYYSFQDKDNLYFVMDYIPGGDLMSLLIKLGIFEEELARFYIAEVTCAVDSVHKMG | 120 |
| sp:LATS2_HUMAN_CT | WVVKLYYSFQDKDSLYFVMDYIPGGDMMSLLIRMEVFPEHLARFYIAELTLAIESVHKMG | 119 |
| sp:LATS1_HUMAN_CT | WVVRLYYSFQDKDNLYFVMDYIPGGDMMSLLIRMGIFPESLARFYIAELTCAVESVHKMG<br>***:*.::****. . **:*:*****:*****: : * * ***** **: * *: ***:.* | 119 |
| EmWarts_CT | FIHRDIKPDNILINRDGHIKLTDFGLCTSFRWTHSSKYWDPVFAIPQPGHEKTTTIPTTT | 179 |
| sp:WARTS_DROME_CT | FIHRDIKPDNILIDRDGHIKLTDFGLCTGFRWTHNSKYYQENGNHSRQDS-----MEPW- | 174 |
| sp:LATS2_HUMAN_CT | FIHRDIKPDNILIDLDGHIKLTDFGLCTGFRWTHNSKYYQK-GSHVRQDS-----MEPS- | 172 |
| sp:LATS1_HUMAN_CT | FIHRDIKPDNILIDRDGHIKLTDFGLCTGFRWTHDSKYYQS-GDHPRQDS-----MDFS-<br>*****: *****:*****:*****:*****:*****:*****:*****: | 172 |
| EmWarts_CT | EAVATAGTTGLAEGDTEGDEGEVGDQVWPMHDQTLERRIRSSSNRRCAKSLVGTPNYIAP | 239 |
| sp:WARTS_DROME_CT | -----EYSE-----NGPKPTVLERRRMRDHQRVLAHSIVGTGPNYIAP | 212 |
| sp:LATS2_HUMAN_CT | -----DLWDDVSNCRCGDRLKTLEQRARKQHQRCLAHSLVGTPNYIAP | 215 |
| sp:LATS1_HUMAN_CT | -----NEWGDPSSCRCGDRLKPLERRAARQHQRCLAHSLVGTPNYIAP<br>. : : **:* . : * *:***** | 215 |
| EmWarts_CT | EILRRQEYNRACDWWSVGVILYEMLVGRPPFVAQTALDTQLRVIQWQRHLRIPGEPRLAS | 299 |
| sp:WARTS_DROME_CT | EVLERSGYTQLCDYWSVGVILYEMLVGQPPFLANSPLETQQKVINWEKTLHIPPQAELSR | 272 |
| sp:LATS2_HUMAN_CT | EVLLRKGYTQLCDWWSVGVILFEMLVGQPPFLAPTPTETQLKVINWENTLHIPAQVKLSP | 275 |
| sp:LATS1_HUMAN_CT | EVLLRTGYTQLCDWWSVGVILFEMLVGQPPFLAQTPLETQMKVINWQTSLHIPPQAKLSP<br>*: * * *. : **:*****:*****:***:* : :** **:*: : **: * : .* | 275 |
| EmWarts_CT | DAADLIRKLLCDPEDRLADP--RLMKQHAFFDSVDWD-KLPTQTPPYVPTILNELDTSNF | 356 |
| sp:WARTS_DROME_CT | EATDLIRRLCASADKRLGKS-VDEVKSHDFFKGIDFA-DMRKQKAPYIPEIKHPTDTSNF | 330 |
| sp:LATS2_HUMAN_CT | EARDLITKLCCSADHRLGRNGADDLKAHPFFSAIDFSSDIRKQPAPYVPTISHPMDTSNF | 335 |
| sp:LATS1_HUMAN_CT | EASDLIIKLCRGPEDRLGKNGADEIKAHPFFKTIDFSSDLRQQSASYIPKITHPTDTSNF<br>: * *** : * . : **. : * * ** . : * : . : * * * : **** | 335 |
| EmWarts_CT | DAVDEEAGGDDQASSQSSCGGAGGGGGTFGLTSAPHAFPDFTFKRFFDQDNNPTTSERDA | 416 |
| sp:WARTS_DROME_CT | DPVDPEKLRSNDSTMSSGD--DV-----DQNDRTFHGFFEFTFRRFFDDKQPPDMTDDQA | 383 |
| sp:LATS2_HUMAN_CT | DPVDEESPWNDASEGSTKA---WDTLTSPNNKHPEHAFYEFTFRRFFDDNGYPFRCPKPS | 392 |
| sp:LATS1_HUMAN_CT | DPVDPDKLWSDDNEEENVN--DTLNGWYKNGKHPEHAFYEFTFRRFFDDNGYPYNYPKPI<br>* ** : . : . . * . * :***:***:. * | 393 |
| EmWarts_CT | I----CSTGAWHQTSAVLVPS-----SSYQ | 437 |
| sp:WARTS_DROME_CT | -----PVYV | 387 |
| sp:LATS2_HUMAN_CT | GAEASQAESSDLESSDLVDQTE-----GCQPVYV | 421 |
| sp:LATS1_HUMAN_CT | EYEYINSQGSEQQSD-EDDQNTGSEIKNRDLVYV | 426 |
|  | * |  |

**Figure S3: Sequence features and homologies of EmWarts.** Depicted is an amino acid sequence comparison between the predicted serine/threonine kinase domains including C-terminal regions of EmWarts (EmuJ\_000700100) and *D. melanogaster* Warts (UniProt Q9VA38) as well as human LATS1 (Uniprot O95835) and LATS2 (Uniprot Q9NRM7). Sites of perfect alignment (\*) as well as groups of strong (:) or weak (.) similarity are marked below the alignment. The predicted serine/threonine kinase domain of EmWarts is marked in blue, the predicted AGC-kinase C-terminal domain is depicted in violet.

Figure S4

|  |  |  |
| --- | --- | --- |
| EmSav | MSRQDSHLFSSKSSGASSVLRPSKLPIFGRRTSEKHPPPTSLTPVSSTVTTSSLGRVRPS | 60 |
| EmSav | TVVRPFFAAQYQPTANNDAQRRRYPASQPSTPVRGGQHETNQEFQQLHRSAHFLRSGGTS | 120 |
| EmSav | GSTTPKRGDIIgaIPQsISfPGSRHALGNQPSKSSKAKSYRMENMSFEHLQYLERRLQQL | 180 |
| EmSav | LEVTSFRPVLSAKAKSDSGRQSTHPSWASSTHHHHHHQHQALHYPVVATPLHYHPHTTPL | 240 |
| EmSav | LLFPQQSIPQPLPTPSLPPPASQLLPPHPTVSGPNFGDIDPYYLSSPSSASSCYAGSSPS | 300 |
| EmSav | SLQDFNRDELICLLHAVRNMLSMHQQSQQSSTSPLDPLSMLQHRPFSWYLHRTADTTAAS | 360 |
| sp: SAV_DROME | --MNY-LTILLC--NRKPTMLSRN-KEKSQ-HKE-----GVVGKYMKK-----DT | 39 |
| sp: SAV1_HUMAN | -----MLSRKKTKEVS-KPA-----EVQGYVKK-----E- | 25 |
|  | *** :: .: . | . *::: |
| EmSav | SPASSTINLFLDEDDSVNVEA-M-----DDGGIVVTDDSKEEESRGR-- | 401 |
| sp: SAV_DROME | PPDISVINVWSDQRAKKKSLQRCASTSPSCEFHPRSSSTSRNTYSCT-DSQPDYYHARRA | 98 |
| sp: SAV1_HUMAN | ----- | 25 |
| EmSav | -----KN-----SASEGGPHPPHLPpGFRNFSRYLDSKFGPDSTEEQRSRQH | 443 |
| sp: SAV_DROME | QSQMPLQQHSHSHPHSLPHPSHPHVRSHPPLPPHQFRASSNQLSQNSSNYVNFE----QI | 154 |
| sp: SAV1_HUMAN | ----- | 25 |
| EmSav | HRHQRRRHREVNDNNTTASVVATTSSSGGNRGRSDTDVRPKISITTCSGGTGPYSDLAPS | 503 |
| sp: SAV_DROME | ERMRR-----QQSSPLLQTTSFA-----PGAGGFQRSYST | 185 |
| sp: SAV1_HUMAN | -----TSPLLRNL----- | 33 |
|  | :: :: . |  |
| EmSav | PHESFLKTTTPNPTSGED-NEIAHRL-APAWCAPYWCP-----PPPLNSGQPTSW | 551 |
| sp: SAV_DROME | T-----QRQHHPHLGGDSYDADQGLLSASYANMLQLPQRPHSPAHYAVPPQQQQHPQIH | 239 |
| sp: SAV1_HUMAN | -----MP----- | 35 |
|  |  | * |
| EmSav | YHHAAPPPPPPL--PPPPPPLPPPQVRYFAPQPPPNAAAACWFRAAANHFATAAAIFAAD | 609 |
| sp: SAV_DROME | QQHASTPFGSTLRFDRaAMSIRERQPRYQ-PTSSPMQOQOQOQOQOQOQLQ--HTQLAAH | 296 |
| sp: SAV1_HUMAN | -----SFIRHGPTIPRRTDICL-PDSSPNAFSTSGDV----- | 66 |
|  | : | * * |
| EmSav | NAAAASVLAAPHQLLPPFPHPSDPTFVDRTREHRAPASS-----SSSTVHPQNVPTT | 661 |
| sp: SAV_DROME | LGGSYSS-----DSYPIYENPSRVIsmRATQSQRSESPiYSNTTASSATLAVVPQHhH-- | 349 |
| sp: SAV1_HUMAN | VSRNQSF-----LRTPIQRTpHEIMR--RESNRLSAPSYLA-----RSLADVPREYG-- | 111 |
|  | . * * * * | ..*: : : : *:: |
| EmSav | SPSNFPSLTLYPPPQFDEGNSIRKGEREEEDEIFLVSDEENQQQGHRQCLRRLPAVEPLQ | 721 |
| sp: SAV_DROME | -----QGHLAVPSGSGGGSLSG-----SG----- | 368 |
| sp: SAV1_HUMAN | SSQSFVTEVSFAVENGDSGSRYYY-----SDNF----- | 139 |
|  | . . * | . |
| EmSav | RQLWGRRSDDSGSPLDSTQTDDDGTWA-----LDMLPQDPALILPD----QVDFYAQIKD | 772 |
| sp: SAV_DROME | -----RGGSSGSVRGASTSV-----QSLY-VPPRTPPSAVAGAGGSANGSLQKVPS | 413 |
| sp: SAV1_HUMAN | -----FDGQRKRPLGDRAHEDYRYYEYNHDLFQRMPQNQGRHASGIGRVAATSLGN--- | 190 |
|  | ... . | *: . : |
| EmSav | LLELDSNISLPLPTGWAVGVSSSGRRFYVSCTTGDAGNGGGGERSISSRNRARTTTWQHP | 832 |
| sp: SAV_DROME | QQSLTEPEELPLPPGWATQYTLHGRKYYIDH-----NAHTTHWNHP | 454 |
| sp: SAV1_HUMAN | -LTNHGSEDLPLPPGWSVDWTMRGRKYYIDH-----NTNTTHWSHP | 230 |
|  | .**** **:. : **:::.. | ..** *.** |
| EmSav | IIAPRIPLGWERVEMRRGNCVYYRHLLIPHTQRHHPDLWFPTSLK----NVEFEQQSWF | 887 |
| sp: SAV_DROME | LEREGLPVGWRRVVSkmHGT-YYENQYTGQSQRQHPCLTSYYVYTTSAEPPKAIRPEASL | 513 |
| sp: SAV1_HUMAN | LEREGLPPGWERVESSEFGT-YYVDHTNKKaQYRHPCAPSVPRYDQP--PPVTYQPQ--- | 284 |
|  | : : * **.* * * . ** . :*: ** | . : |
| EmSav | FDL-RKLQESVS---NFDKGNSQLIESYVDARDSNEEAHFTALLKQHdVGQLEKLLKHL | 942 |
| sp: SAV_DROME | YAPPTHTHNALVPANPYLLEEIPKWLAVYSEADSSKDHLLQFNMF---SLPELEGfDSML | 570 |
| sp: SAV1_HUMAN | --QTERNQSLlVPANPYHTaEIPDWLQVYARAPVKYDHILKWELF---QLADLDTYQGMl | 339 |
|  | : : . : . : * * . . : : :*: * |  |
| EmSav | DCRFFRDLHRMVVAfEVARIRIVRELFVQHIHRTTATTASSPpSAIHGSAEK | 994 |
| sp: SAV_DROME | VRLFKQELGTIVGFYERYRRALILEKNR---RAGQNQ---NQNQ----- | 608 |
| sp: SAV1_HUMAN | KLLFMKELEQIVKMYeAYRQALLTELEN---RKQRQq---WYAQQHGKNf- | 383 |
|  | * :*: * : * * : : * | * |

**Figure S4: Sequence features and homologies of EmSalvador.** Depicted is an amino acid sequence comparison between full length EmSalvador (EmSav; EmuJ\_000331400) and Sav orthologs from *D. melanogaster* (UniProt Q9VCR6) and *H. sapiens* (Uniprot Q9H4B6). Sites of perfect alignment (\*) as well as groups of strong (:) or weak (.) similarity are marked below the alignment. A predicted 7BQF domain typical of Sav proteins is indicated in blue.

**Figure S5: Sequence features and homologies of EmScalloped.** Depicted is an amino acid sequence comparison between full length EmScalloped (EmScal; EmuJ\_000606300) and Scalloped orthologs from *D. melanogaster* (UniProt P30052) and *H. sapiens* (Uniprot P28347). Sites of perfect alignment (\*) as well as groups of strong (:) or weak (.) similarity are marked below the alignment. A predicted TEA transcription factor domain of EmScal is indicated in blue, a 5Z2Q PDB domain typically found in Scalloped proteins is marked in violet.

|  |  |  |
| --- | --- | --- |
| EmSca1 | MQPPASSGSRMPLPGQSNPSVRAPPSSSTTAHAGLPSPFDYPPMHNMVHGFRSPQQPPFK | 60 |
| sp: SCAL_DROME | -----MK | 2 |
| sp: TEAD1_HUMAN | ----- | 0 |
| EmSca1 | EVASPAAALPSAYDMAAVFISASSSAAAAAAAASAASLRPFGPSAQKRPATISPLTLGSG | 120 |
| sp: SCAL_DROME | NITSSSTCSTGLLQLQNNL-S--C---SELEVAEKTEQQAVGPGT--IPSPWTPVNAGPP | 54 |
| sp: TEAD1_HUMAN | -----M-EPSSWSGSE---- | 10 |
|  | *: : |  |
| EmSca1 | GGGGGGSSPVITGGSECDATTASQSDDAEVDGGAGGGGGGNEADGVWSTEIEQYFQEALE | 180 |
| sp: SCAL_DROME | GALG----SADTNGSMVDS----KNLDVGDMSDDEKDLSSADAEGVWSPDIEQSFQEALS | 106 |
| sp: TEAD1_HUMAN | -----S----PAENMERMSDSADKPIDNDAEGVWSPDIEQSFQEALA | 48 |
|  | : : . . :*:*** :*** ***** |  |
| EmSca1 | LYPPCGRRKIILAEEGKMYGRNELIARYIQQRTGKVRTRKQVSSHIQVLARRKSKELQAK | 240 |
| sp: SCAL_DROME | IYPPCGRRKIILSDEGKMYGRNELIARYIKLRTGKTRTRKQVSSHIQVLARRKLREIQAK | 166 |
| sp: TEAD1_HUMAN | IYPPCGRRKIILSDEGKMYGRNELIARYIKLRTGKTRTRKQVSSHIQVLARRKSRDFHSK | 108 |
|  | :*****::*****: ****.***** :::: |  |
| EmSca1 | IRDPEIKKETINQLAKLSSAQIVSRDVAKGLQSEGNVKFLPQQPLPSPTSSCSSTAAAAA | 300 |
| sp: SCAL_DROME | IKVQF----- | 171 |
| sp: TEAD1_HUMAN | LKDQTAKDKALQHMAAMSSAQIVSATAIHNK----- | 139 |
|  | :: |  |
| EmSca1 | ATVAASNQLHHAFHGQQQHHHQHQRQQVPYRQAHATRKSSIGEGASQRKMMRLADPEVYN | 360 |
| sp: SCAL_DROME | ----- | 171 |
| sp: TEAD1_HUMAN | ----- | 139 |
| EmSca1 | TTDSFLSLNNGQNPAVHAARSEDLKLCSSSISQRVVMSGPPPNAFPPDTHYSSLYVRSHN | 420 |
| sp: SCAL_DROME | -----W----- | 172 |
| sp: TEAD1_HUMAN | -----LGLPGIPRPTFPGAPGFW----- | 157 |
| EmSca1 | PNPHHNPHLAQHTTPLPPLLEMGGCSSDRCVSSAAPPPLQHPPPPPPPPPPDFSAMQLAA | 480 |
| sp: SCAL_DROME | -----QPGLQPSTS-QDF--YDISIKFPFPYPYPAG--KTS----- | 203 |
| sp: TEAD1_HUMAN | -----PGMIQTG---QPG--SSQDVKFVFQYPIQ---PAV----- | 186 |
|  | : : . . * : : * |  |
| EmSca1 | AAGLEHYFSSGRSSSIDTCLYPNAFQPPPPPLPPPPPPPPPSSQSVALTAAPLPLSDFDM | 540 |
| sp: SCAL_DROME | -----TA-VSG--DETG-----IPPSQ | 217 |
| sp: TEAD1_HUMAN | -----TAPIPGFEPASA-----PAPSV | 203 |
| EmSca1 | QWWINRSIASDKLRLVELCAFVEYPCQISECAAEDLTLNPESAPMKQHNFAHITTTGLHS | 600 |
| sp: SCAL_DROME | LPWEGRAIATHKFRLLEFTAFMEIQRD-----EIYHRHLFVQLGG-KPSF | 261 |
| sp: TEAD1_HUMAN | PAWQGRSIGTTKLRLVEFSAFLEQQRD-----PDSYNKHLFVHIGHANHSY | 249 |
|  | * .*:.: *:***: ***: : :*: *:: |  |
| EmSca1 | SDPILEEVDASQIWDKFPT--DGLKQRIEEDSPNVFFLVKLWADINTELPE----GANYA | 654 |
| sp: SCAL_DROME | SDPLLETVDIRQIFDKFPEKSGGLKDLYEKGPQNAFYLVKCWADLNTDLTTGSETGDFYG | 321 |
| sp: TEAD1_HUMAN | SDPLLESVDIRQIYDKFPEKKGGLKELFGKGPQNAFFLVKFWADLNCNIQ--DDAGAFYG | 307 |
|  | ***:* ** **:**** .***: :. *.*** ***: * : : * |  |
| EmSca1 | VSSIFEGTEDVPLLVSTRLCSYSSPFVEKLEYESPVLENGRYVYKQLRSPMCACMKKFIA | 714 |
| sp: SCAL_DROME | VTSQYESNENVVLVCSTIVCSFGKQVVEKVESEYSRLENNRYVYRIQRSPMCEYMINFIQ | 381 |
| sp: TEAD1_HUMAN | VTSQYESSENMTVTCSTKVCSFGKQVVEKVETEYARFENGRFVYRINRSPMCEYMINFIH | 367 |
|  | *:* :*.:.: : ** :***.. .***:* * :**.*:*** :***** * :* |  |
| EmSca1 | KLLKLSSLAE MNQVLENFTILQIVTNKNTDEVLLGIAYVLEVSKSLRG AQHNIYRLAKPD | 774 |
| sp: SCAL_DROME | KLKNLPERYMMNSVLENFTILQVMRARETQETLLCIAYVFEVAAQNSGTTHHIYRLIKE- | 440 |
| sp: TEAD1_HUMAN | KLKHLPEKYMMNSVLENFTILLVVTNRDTQETLLCMACVFEVSNSEHGAQHHIYRLVKD- | 426 |
|  | ** :* . **.****** : : :*:.* ** :* :*: . * :* ***** * |  |
| EmSca1 | ISSTNQRPPSVNLNGQPLPATSTVD | 799 |
| sp: SCAL_DROME | ----- | 440 |
| sp: TEAD1_HUMAN | ----- | 426 |

Figure S6

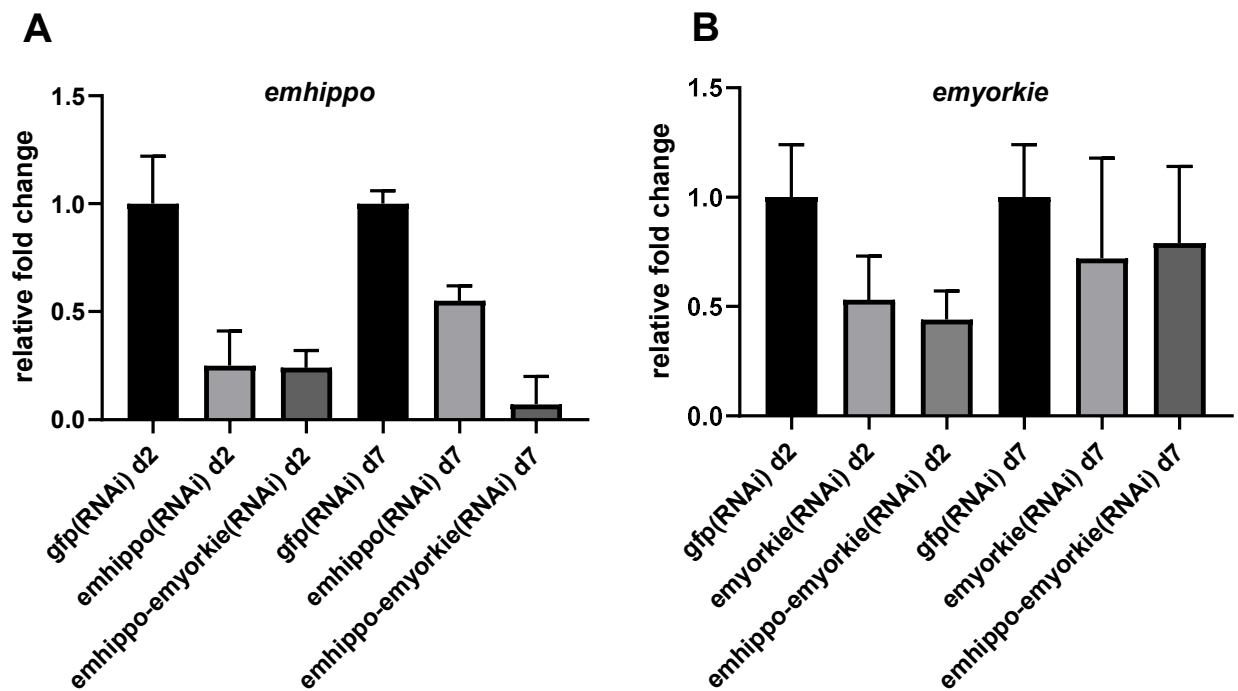

**Figure S6. RT-qPCR analysis of *emhippo* and *emyorkie* expression following RNAi treatment.** Relative transcript levels of *emhippo* (A) and *emyorkie* (B) were determined by RT-qPCR at 2 and 7 days after initiation of RNAi treatment. Primary cell cultures were subjected to RNAi targeting *emhippo*, *emyorkie*, or both genes simultaneously, as indicated. Transcript levels were normalized to those of the corresponding GFP-RNAi controls, which were set to 1 for each time point. At day 2, reduced transcript levels were detected for both target genes following the respective RNAi treatments. At day 7, *emhippo* expression remained reduced, whereas *emyorkie* transcript levels showed partial recovery. Bars represent mean values, and error bars indicate the standard deviation (SD) from technical RT-qPCR measurements of one biological experiment.

Figure S7

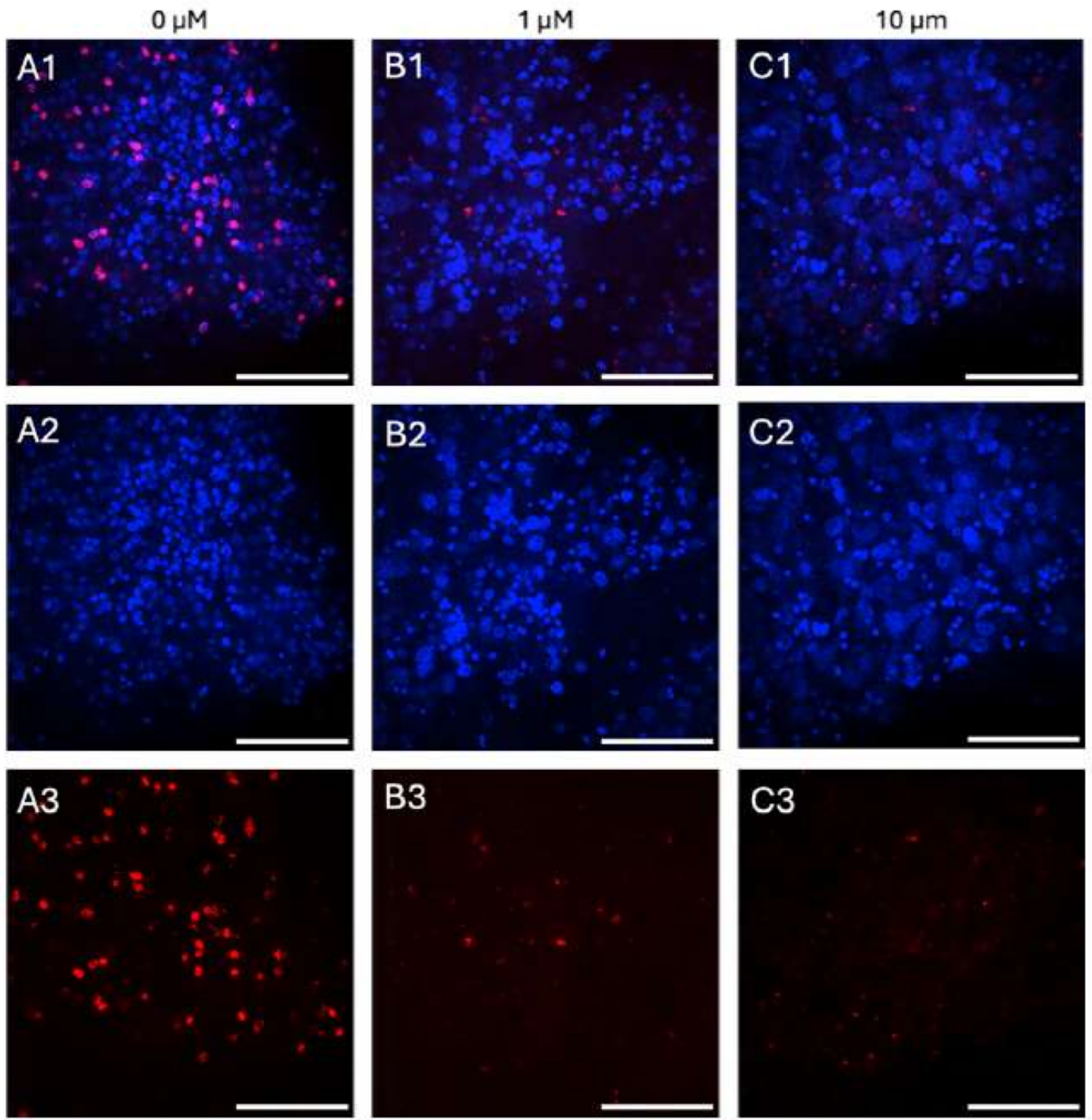

**Figure S7. XMU-MP-1 treatment reduces cell proliferation and induces morphological alterations in *E. multilocularis* primary cell cultures.**

Representative fluorescence microscopy images of primary cell aggregates treated with 0 μM (A1–A3), 1 μM (B1–B3), or 10 μM (C1–C3) XMU-MP-1. Merged images (A1–C1) show nuclear staining with DAPI (blue) and EdU incorporation (red), indicating proliferating cells. The individual DAPI (A2–C2) and EdU (A3–C3) channels are shown below. XMU-MP-1 treatment resulted in a pronounced reduction of EdU-positive cells at both concentrations, accompanied by alterations in cell morphology, including the appearance of enlarged cells, particularly at 10 μM. Scale bars as indicated.

**Figure S8**

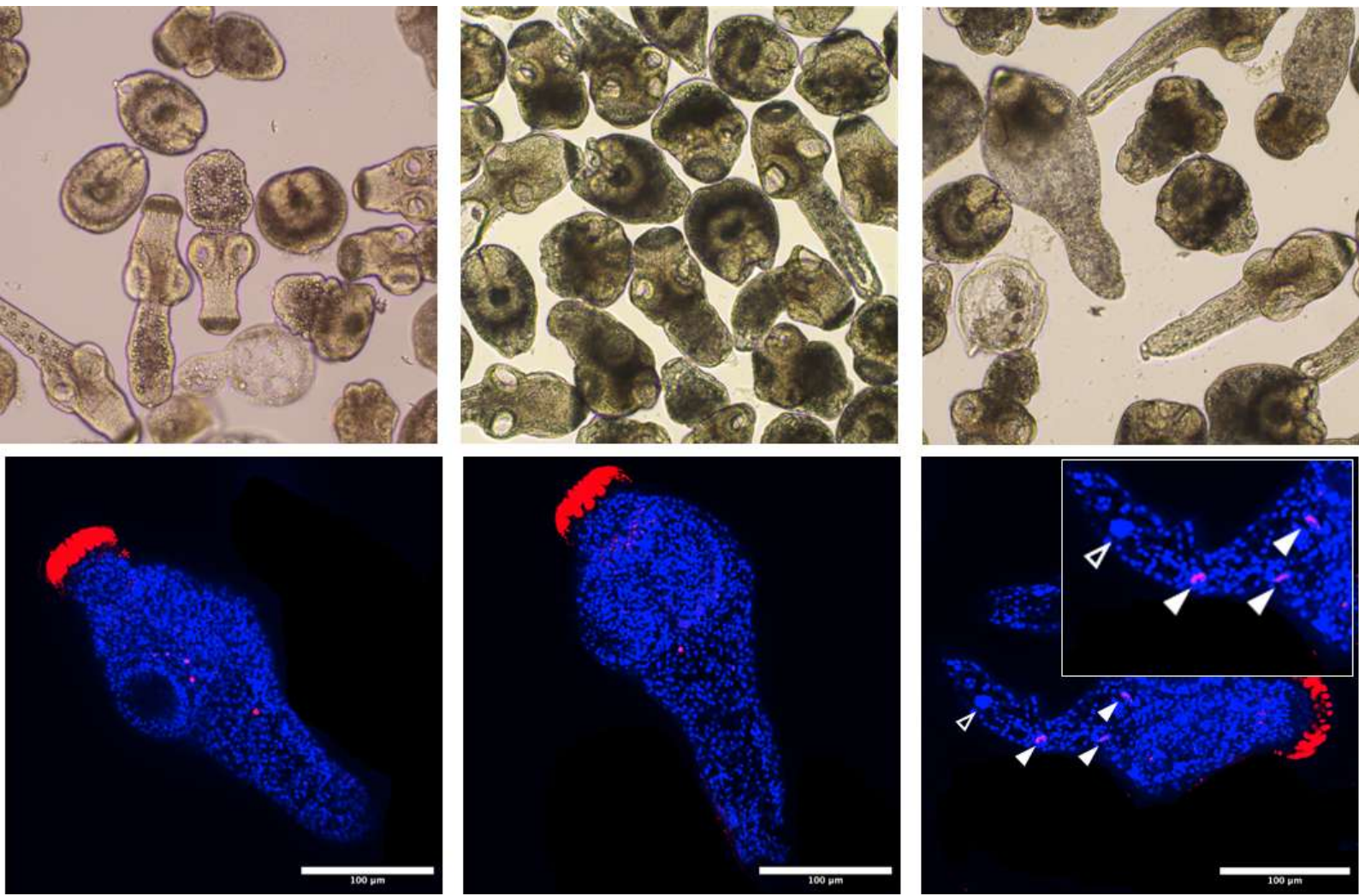

**Figure S8. XMU-MP-1 treatment induces morphological alterations and enlarged cells in *E. multilocularis* protoscoleces.**

Representative bright-field (upper panels) and fluorescence microscopy images (lower panels) of *E. multilocularis* protoscoleces following treatment with 0 $\mu\text{M}$ (left), 1 $\mu\text{M}$ (middle), or 10 $\mu\text{M}$ (right) XMU-MP-1 for 14 days. Nuclei were stained with DAPI (blue), while EdU incorporation (red) indicates proliferating cells. Treatment with XMU-MP-1 induced concentration-dependent morphological alterations, including the appearance of enlarged cells at 10 $\mu\text{M}$. Arrowheads indicate enlarged cells within the protoscolex tissue, shown at higher magnification in the inset. Scale bars: 100 $\mu\text{m}$ (fluorescence images).

Figure S9

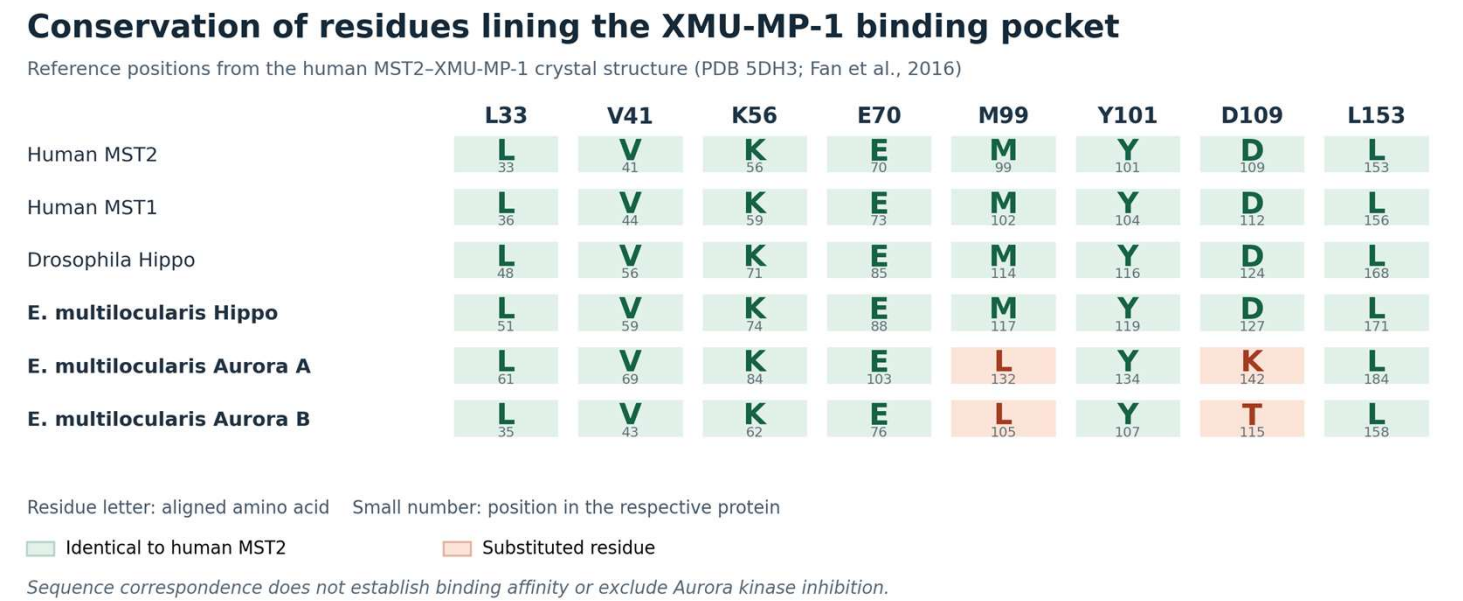

**Figure S9. Conservation of amino acid residues implicated in XMU-MP-1 binding among Hippo and Aurora kinases.** Comparison of eight amino acid positions lining the XMU-MP-1-binding pocket of human MST2, as identified by structural analysis (Fan et al., 2016), with the corresponding positions in human MST1, *Drosophila* Hippo, and *Echinococcus multilocularis* Hippo, Aurora A, and Aurora B kinases. Residues identical to human MST2 are highlighted in green, whereas substitutions are indicated in orange. All eight positions are conserved in EmHippo, whereas both EmAURKA and EmAURKB exhibit substitutions at positions corresponding to MST2 M99 and D109. The Aurora kinase sequences were derived from Cheng et al. (2019). These sequence differences suggest potential alterations in inhibitor recognition but do not establish differences in binding affinity or kinase inhibition.
